# Ovary-Derived Signals Align Protein Appetite with Oogenesis

**DOI:** 10.64898/2026.03.23.711327

**Authors:** R. R. Nóbrega, A. P. Francisco, A. M. Gontijo, C Ribeiro, Z. Carvalho-Santos

## Abstract

Maintaining organismal homeostasis requires mechanisms that coordinate the metabolic needs of individual organs with whole animal nutrient intake. Although nutrient sensors and central pathways regulating hunger have been extensively characterized, how peripheral organ physiology influences nutrient specific appetites remains poorly understood. Here, we identify a previously unrecognized signalling axis, originated in the ovary, that modulates yeast appetite in *Drosophila melanogaster*. Through a targeted germline RNAi screen, we find that specific perturbations in oogenesis consistently and selectively increase yeast appetite. The manipulations increasing yeast appetite disrupt oogenesis progression, producing a shared signature of increased vitellogenic follicle accumulation and reduced number of mature (stage14) oocytes. This shift is accompanied by decreased expression of the relaxin-like hormone Dilp8, and loss of Dilp8 recapitulates the feeding phenotype. We show that this ovarian regulation of nutrient specific appetite is independent of amino acid state but requires mating, indicating integration with Sex Peptide–mediated reproductive activation. Together, our findings uncover a novel mechanism that couples oogenesis progression to nutrient selection, emphasizing the importance of the ovary as an active regulator of whole organism nutritional decisions. This work provides a conceptual framework for how reproductive tissues communicate their physiological demands to other organs and raises the possibility that analogous ovary-derived signals may shape nutrient specific appetite and metabolic states in other animals.

**Graphical Abstract:** 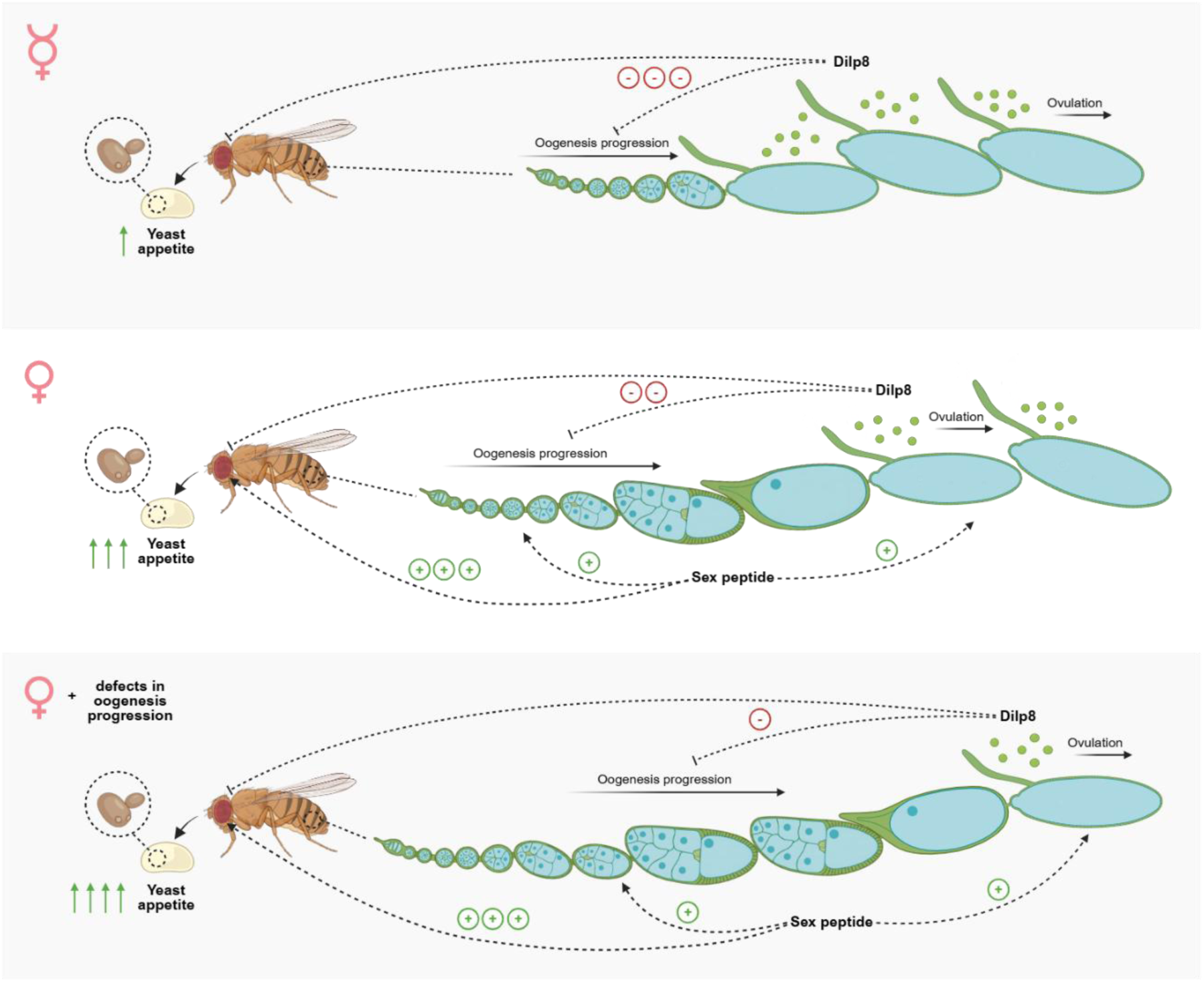

Created in BioRender. Francisco, A. (2026) https://BioRender.com/pwalhtl

## Introduction

Multicellular animals are composed of diverse organs, each containing specialized tissues with unique demands. A fundamental challenge for whole-animal physiology is the coordination of these distinct requirements, ensuring tissue functions, while preserving homeostasis. To achieve this, animals rely on complex inter-organ communication networks, mediated by secreted metabolites, proteins, and peptides that convey the physiological status of individual organs, allowing systemic coordination^1–5^. These signaling pathways play a central role in mediating the responses to internal fluctuations or external perturbations. Work carried out across multiple model organisms, including mouse, *Caenorhabditis elegans* and *Drosophila melanogaster*, has significantly advanced our understanding of how organs exchange information to regulate processes such as development, reproduction, energy homeostasis, or immune function^1–5^. However, despite this progress, we still do not fully understand how organs sense and coordinate each other’s needs, and how the integration of such information shapes whole-animal physiology and behavior.

Animals continuously monitor numerous internal variables, including temperature, circulating gases, and accumulation of toxic metabolites, to maintain organ functions within optimal ranges. Among these variables, nutrient homeostasis emerges as one critical regulator of animal physiology^6,7^. The amount and composition of ingested food determine the availability of specific metabolites to individual tissues, influencing their function. As a consequence, dietary composition and quantity have profound effects on whole-animal performance and life-history traits such as fecundity, lifespan, and aging^8–19^. Although nutrient availability and balance clearly trigger specific physiological programs, how individual nutrient classes are sensed, how their availability is communicated across organs, is still poorly understood.

Given the strong influence of nutrition on physiology, animals have evolved sophisticated mechanisms to adjust feeding behavior according to internal needs, ensuring adequate intake of nutrients and calories while preventing excessive consumption, a process broadly referred to as homeostatic regulation of food intake^2–4,6,20–23^. A substantial body of work has focused on general “hunger” and “satiety” signals and on the neuronal mechanisms that mediate the physiological and behavioral responses downstream of these signals^22–26^. However, how appetites for specific nutrients are regulated to achieve nutritional homeostasis and optimize fitness remains less clear. Classic behavioral studies demonstrated that animals can selectively regulate the intake of individual macro- and micronutrients^27–33^. In *Drosophila*, nutrient-specific appetites constitute a key component of this regulation: deprivation of carbohydrates or amino acids elicits a selective increase in the consumption of sugar- or protein-rich foods, respectively^6,8,34–39^. Feeding behavior can also be adjusted in anticipation of future physiological demands. For instance, mating induces female flies to increase their appetite for protein and salt, anticipating the energetic and biosynthetic requirements of egg production^6,37–40^. Despite these advances, the molecular and cellular mechanisms by which dietary information is integrated with tissue-specific physiological demands to regulate nutrient-specific appetite remain incompletely understood.

In *Drosophila,* female fertility is highly sensitive to dietary composition, with the ovary responding dramatically and readily to various dietary challenges including starvation, dietary restriction, the complete removal of specific nutrients, and shifts in macronutrient ratios^8,17,34,36,41–51^. In *Drosophila*, oocytes are produced from a series of steps involving both germline and somatic tissues within the ovary. The germline stem cells, located at the anterior tip of the germarium, divide to produce one cystoblast while the other cell remains in the niche. The cystoblast undergoes four rounds of mitotic divisions, after which one cell is specified into the oocyte while the remaining 15 will become nurse cells. These 16 germline cells are surrounded by somatic tissue, the follicular epithelium, forming an egg chamber that detach from the germarium to undergo 14 developmental stages until it becomes a fully formed oocyte^52^. Protein-poor diets lead to a dramatic reduction of egg production resulting from a combination of effects at the level of oogenesis, including a reduction of germline and follicle stem cell proliferation and maintenance, a block of vitellogenesis through decreased uptake of yolk and lipids into the oocyte, and an increase in autophagy and apoptosis of mid-oogenesis egg chambers^42,53–57^. Because ovarian function is essential for animal fitness, nutrient intake should be tightly aligned with its metabolic demands. Yet, how the ovary communicates its nutritional needs to the rest of the animal ensuring a continuous supply of essential nutrients is yet to be fully elucidated.

Consistent with this idea, recent work has begun to reveal that, in *Drosophila*, nutritional state is not the only modulator of feeding behavior but that the physiological state of peripheral organs, namely the ovary, can influence animal appetite. For instance, ovarian carbohydrate metabolism regulates sugar appetite: the pentose phosphate pathway in the germline is essential for egg production, and its disruption reduces sugar intake independently of circulating sugars^36^. Manipulating germline metabolism affects expression of the fat-body-derived satiety factor FIT, which acts through the central nervous system to modulate feeding^36,58^. These findings suggest that the physiological state of the ovary is actively monitored to tune nutrient-specific appetites. Despite this evidence, the mechanisms by which ovarian physiology is monitored, how this information is encoded and transmitted, and how it modulates nutrient-specific appetites is still far from fully understood.

In this study, we uncovered an ovary-mediated mechanism that modulates yeast-specific appetite in *D. melanogaster*. Using a targeted RNAi screen in the female germline, we identified a set of genes whose knockdown selectively increases yeast appetite without affecting sugar intake. These manipulations disrupt oogenesis progression, consistently reducing the number of stage 14 oocytes. We show that these changes in oogenesis are linked to a decrease in expression of the relaxin-like hormone Dilp8, whose knockdown phenocopies the feeding effects. Our data indicates the existence of ovary-derived signals that reports the progression status of oogenesis and modulates yeast appetite in a mating-dependent but amino acid-state independent manner. Together, our findings reveal a previously unrecognized endocrine axis linking oocyte development to central regulation of nutrient-specific appetite, demonstrating how reproductive physiology contributes to behavioral homeostasis.

## Materials and Methods

### Drosophila melanogaster stocks and genetics

Germline expression of transgenes for RNA interference was achieved using MTD-Gal4 (Maternal Triple Driver) unless otherwise stated. In specific experiments, the maternal driver *matα4*-Gal4 was used. Follicle cell expression of transgenes for RNA interference was achieved using *tj*-Gal4 (Gontijo Lab). shRNA lines used in this study were obtained from the TRiP collection ^59,60^. The corresponding controls were chosen according to the vector backbone where shRNA was originally cloned and the insertion site of the experimental shRNA lines. *dilp8* lhRNA and Green Fluorescence Protein (GFP) 30B lhRNA transgenic lines were obtained from the Vienna Drosophila Resource Center (VDRC)^61^. The *dilp8* knockout mutant (*dilp8^ag^*^50^) and respective control (*dilp8^ag^*^52^) were generated by Alisson Gontijo^62^. Drosophila Genetic Reference Panel (DGRP)-426 was used as wild type^63^. Information regarding all stocks used in this study is detailed in **Supplementary Table 1.**

We used FlyBase (FB2024 and FB2025 releases) to retrieve information on gene-associated phenotypes, functions, and stocks^64,65^.

### Drosophila melanogaster rearing, media, and dietary treatments

Flies were reared in Yeast Based Medium (YBM) until adulthood (see **Supplementary Table 2** for diet detailed composition). Holidic Media (HM) were prepared as described previously, using the FLYaa formulation^8,17^. The different HM used in this study are described in **Supplementary Table 3**.

To ensure a homogeneous density of offspring among experiments, fly crosses were always generated in YBM with six females and three males per vial. In all experiments, drivers were carried by females and RNAi transgenes by the males. The following dietary treatment protocol was used to ensure a well-fed and mated state: groups of one- to four-day-old flies (16 females and 5 wild type (WT) males for behavioral assays or 8 females and 4 WT males for fertility assays) were selected from the crosses, 14 days after cross setup, and collected into fresh YBM-containing vials. Unless otherwise stated, flies were always maintained for 48 hours in YBM and transferred into a complete HM for four days before assays were performed. This protocol enabled targeted dietary manipulations while ensuring consistency of nutritional conditions across all experimental paradigms described in this manuscript.

For assessing the effects of nutrient specific deprivation, flies were transferred to the correspondent HM (**Supplementary Table 3**) for 4 days before fertility and behavioral assessment.

For the mating status experiments, virgin females were collected for four days, starting from 10 days after cross setup. For the mated conditions, groups of 16 virgin females were transferred to fresh vials of YBM and housed with 5 males. For the virgin conditions, groups of 21 virgin females were transferred to fresh vials of YBM. Females were kept under these conditions for 6 days before the behavioral assays.

Flies were always reared at 25°C, 70% relative humidity and on a 12-hour light-dark cycle.

### Egg-laying assays

Groups of 8 female and 4 male flies were briefly anesthetized using light CO2 exposure and transferred to apple juice agar plates (composition described in **Supplementary Table 4**), where they were allowed to lay eggs for 24 hours. Parents were then removed and females counted and the plates placed at 25°C for 48 hours to ensure eggs had sufficient time to hatch. The number and viability of eggs was then assessed. Number of eggs per female was calculated by dividing the total number of eggs in a plate by the number of living females after the 24-hour period. Fraction of viable eggs was calculated by dividing the number of empty eggshells by the total number of eggs. Estimated progeny was calculated by multiplying the average number of eggs per female by the fraction of viable eggs.

### Fly Proboscis Activity Detector (flyPAD) assays

flyPAD assays were performed as described by Itskov and colleagues^35^. Single flies were tested in arenas that contained two food patches: 10% yeast and 20mM sucrose each mixed with 1% agarose. Flies were individually transferred to flyPAD arenas by mouth aspiration and allowed to feed for 1h at 25 °C and 70% relative humidity. The total number of sips per animal during the assay and other feeding parameters were calculated using previously described algorithms^35^. Flies that did not eat (defined as having fewer than two activity bouts during the assay) were excluded from the analysis.

### Colorimetric feeding assays

Food intake was quantified using a colorimetric ingestion assay based on a non-absorbable blue dye (Erioglaucine disodium salt, 1% w/v), as described by Deshpande and colleagues^66^. Dye-containing holidic media (HM) was distributed onto Petri dishes by pipetting nine 10 µL droplets evenly spaced per dish. For each assay, groups of pre-sorted 16 females and 5 males were anesthetized with CO2, transferred to the dish and allowed to feed for 2 h at 25 °C and 70% humidity. Plates were then frozen at –20°C for at least 30 minutes to terminate feeding and immobilize flies.

For quantification, around 30 flies were homogenized in 150 µL of 1% Triton X-100 in PBS (PBS-T) containing a metal bead using a Tissue Lyser (30 Hz, 2 minutes). Homogenates were then centrifuged at 13,300 rpm for 10 minutes, and 40 µL of the supernatant was transferred to a 96-well plate. A standard curve was prepared from serial dilutions of the dyed food solution (initial concentration = 0.01 g/mL). Absorbance was measured at 630 nm using a TECAN Spark microplate reader. Absorbance readings were normalized to the blank control, and dye concentration was calculated from the standard curve to estimate relative food intake per sample. Dye concentration was normalized for the number of females per sample.

### Ovary dissection, staining, and imaging

Ovaries were dissected on ice-cold PBS 1x and fixed with a solution of 4% paraformaldehyde (PFA; 158127, Sigma-Aldrich) and 0.1% Triton X-100 (21123, Sigma-Aldrich) in PBS 1x for 20 minutes at room temperature using soft agitation. Ovaries were washed two times with 0.1% Triton X-100 in PBS (PBS-T).

To quantify egg chamber stages, after fixation and washings with PBS-T, ovaries were incubated with Phalloidin (A30107, Invitrogen) for one hour and then washed twice with PBS-T and once with PBS 1x, before mounting in VECTASHIELD (H-1000, Vector Laboratories). Analysis of the tissue and image acquisition was carried out using ZEISS Axioscan 7 with a Plan-Apochromat 20×/0.8 objective lens and processed using Fiji (ImageJ 1.52n)^67^ and Adobe Photoshop CS5.5. Quantification was performed manually using the morphology of the egg chambers for staging according to Jia and colleagues^68^.

For whole ovary imaging, after fixation and washings with PBS-T, ovaries were incubated with DAPI (62248, Thermo Scientific) for 45 minutes and then washed twice with PBS-T and once with PBS 1x, before mounting in VECTASHIELD (H-1000, Vector Laboratories). Image acquisition was carried out using a Zeiss LSM 980-32 with Airyscan 2. Samples were imaged with a Plan-Apochromat 20×/0.8 objective lens. Following acquisition, individual image tiles were stitched together in ZEN using the built-in stitching module to generate composite images and processed using Fiji (ImageJ 1.52n)^67^ and Adobe Photoshop CS5.5.

Microscopy experiments were performed at the Bioimaging Platform of the Gulbenkian Institute for Molecular Medicine.

### RNA Extraction and cDNA Synthesis

For *dilp8* mRNA quantification, total mRNA was extracted from the fly’s ovaries (eight flies per condition). Tissues were dissected in PBS 1x (diluted in RNAase free water (AM9906, Invitrogen) and snap-frozen in dry ice and 1 mL of purezol (7326890, Bio-Rad) was added before being stored at -80°C overnight. Samples were then thawed at RT for 5 minutes, and the tissue homogenized using a Tissue Lyser (25 Hz, 2 minutes (50985300, Qiagen)) using metallic beads. Subsequently, 200 µl chloroform (288306, Sigma) was added to the samples followed by 15 seconds of vigorous shaking and finally samples were centrifuged at 12 000g for 15 minutes at 4°C. The supernatant was transferred to a new Eppendorf, and 70% cold ethanol was added in the same proportion. The lysate was then loaded into an NZYSpin Binding Column (MB13402, NZYtech), with column purification, washing, and elution being performed according to the manufacturer’s instructions. After RNA quantification using a NanoDrop 2000 spectrophotometer (Thermo Scientific), all samples were diluted to the lowest concentration registered. Complementary DNA (cDNA) was synthesized from 1 µg of RNA using a reverse transcription reaction (2 µL of Buffer, 1 µL of dNTPs (MB08603, NZYtech), 1 µL of random hexamers (MB12901, NZYtech), 0.5 µL of RNA inhibitors (MB08401, NZYtech), and 1 µL of RNA transcriptase (MB12401, NZYtech)). The reaction was performed in a thermal cycler (Bio-Rad) with an initial incubation at 25°C for 10 minutes to allow primer annealing, followed by 55°C for 30 minutes for cDNA synthesis. The reverse transcriptase enzyme was subsequently inactivated at 85°C for 5 minutes. The reaction was then held at 4°C until further processing. The synthesized cDNA was stored at −20°C.

### Quantitative Real-Time PCR (RT-qPCR)

*dilp8* gene expression was determined using real-time PCR. Each cDNA sample was amplified using RT-PCR mix on the Viia7 real-time PCR system (Applied Biosystems). Briefly, the reaction conditions consisted of 1 µg cDNA, 5 µL of Sybr Green (1725124. Bio-Rad), 0.2 µL of each primer and 3.6 µL of water. The cycling program consisted of an initial hold stage at 50°C for 2 minutes and 95°C for 10 minutes. This was followed by 39 cycles of amplification at 95°C for 15 seconds and 60°C for 1 minute. A melt curve analysis was subsequently performed at 95°C for 15 seconds, 60°C for 1 minute, and 95°C for 15 seconds to assess amplification specificity. The primers used in this reaction are listed in **Supplementary Table 5**. Five experimental replicates and two technical replicates per genotype were used. Relative levels of RNA were calculated from a standard curve and normalized to two internal controls (*Act42A* and *Rp49*). The relative quantification of each mRNA was performed using the delta Ct method.

### Data analysis

Effect sizes were quantified using Cohen’s *d.* For each gene and its matched control, the mean and standard deviation was calculated. Cohen’s *d* was calculated using the standard independent-samples formulation. A pooled standard deviation was first computed to estimate the shared dispersion of the condition and control groups:

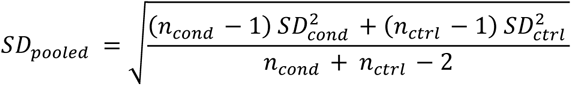

The effect size was then obtained as:

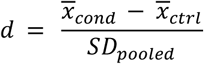

Positive values of *d* indicate an increase relative to control and negative values indicate a decrease. Cohen’s d was only computed when both groups contained sufficient replicates to define a standard deviation; for conditions with a single replicate, Cohen’s *d* is not reported. Values around ±0.8 or greater were considered as large effects.

To calculate correlations between parameters across conditions, median values were first determined for each condition and its corresponding control group. For each parameter, the median of the control group was subtracted from the median of the respective experimental condition to obtain the change relative to control (Δ median). This resulted in one Δ median value per condition for each measured parameter.

To assess correlation between two different parameters, the Δ median values from multiple experimental conditions were plotted against each other in scatter plots. Each data point represented a single experimental condition, defined by its Δ median values for both parameters. A simple linear regression analysis was then performed to evaluate correlation between the two parameters. A linear regression line was fitted to the data, and the strength and direction of the correlation were determined based on the slope and the coefficient of determination (R²). Statistical significance of the correlation was assessed using the p value obtained from the linear regression analysis.

### Statistical Analysis

All statistical analyses were performed using GraphPad Prism 8.4.3. Data was first screened for outliers using the Robust Regression and Outlier Removal (ROUT) method, with a false discovery rate (Q) set at 1%. Identified outliers were excluded prior to further analysis.

Data distribution was assessed for normality using the normality tests available in Prism. The choice of statistical tests was based on the outcome of the normality assessment. For datasets that followed a normal distribution, comparisons between two groups were performed using an unpaired Student’s t-test, and comparisons among three or more groups were conducted using ordinary one-way analysis of variance (ANOVA) followed by Dunnett’s multiple comparisons test. For all other datasets, nonparametric tests were applied: the Mann–Whitney test for comparisons between two groups and the Kruskal–Wallis test followed by Dunn’s multiple comparisons test for comparisons among three or more groups. All tests were two-tailed, and a p value < 0.05 was considered statistically significant. Data are presented as mean ± standard deviation (SD) unless otherwise specified.

## Results

### Ovarian physiology specifically modulates yeast appetite

Given the high nutritional demands of oogenesis and the central contribution of female fertility to organismal fitness, we hypothesized that oogenesis progression must be monitored at the whole-animal level, allowing nutrient intake to be adapted to the ovary’s requirements. If this is the case, perturbing ovarian physiology should alter nutrient-specific appetites.

To test this, we performed a small-scale screen in which we knocked down the expression of 45 genes previously reported to cause embryonic lethality^69^. We reasoned that these manipulations would disrupt germline physiology without fully ablating it and thereby used them as a toolkit to modulate oogenesis. Gene knockdown was restricted to the female germline using MTD-Gal4 (Maternal Triple Driver-Gal4), a strong germline-specific driver, combined with shRNA lines previously linked to fertility defects, or GFP shRNA as a negative control (**Fig. 1a**). We first characterized the impact of these manipulations on ovarian function by measuring female fertility, quantified by the number of eggs laid and their viability, relative to the corresponding genetic controls (**Fig. 1a-b**). As expected, most knockdowns impaired fertility, either by reducing egg production, decreasing egg viability, or both (**Fig. 1b-c** and **Supplementary Fig. 1a-b**). Overall, the estimated progeny was reduced in 26 knockdowns (**Fig. 1d** and **Supplementary Fig. 1c**).

**Figure 1.**
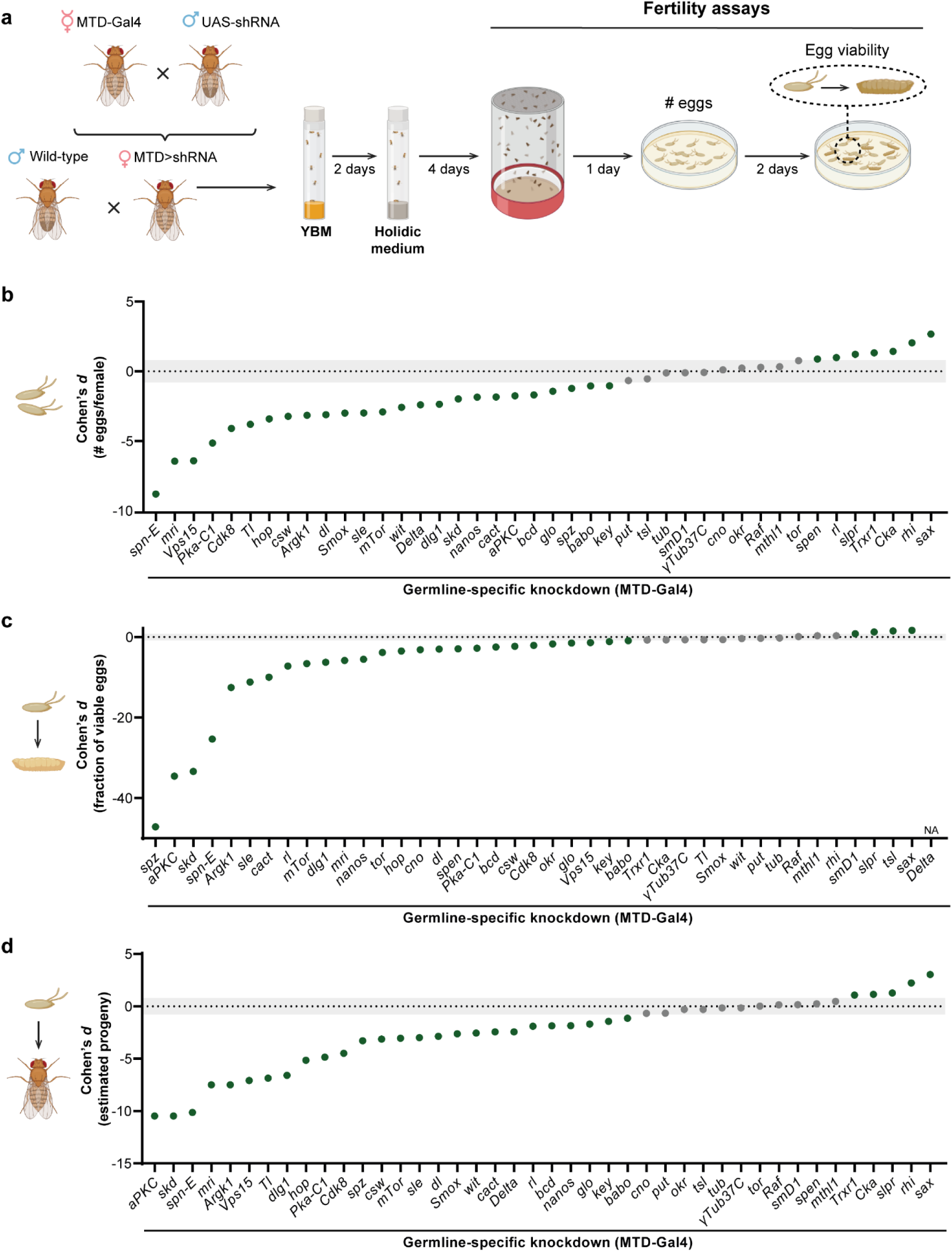
Germline-specific gene knockdowns lead to decreased female fertility. **a** Schematic representation of experimental design and fertility assays. Further details can be found in the Methods section (YBM-Yeast Based Medium). **b**-**d** Fully fed mated females knocked down for the indicated genes using the MTD-Gal4 driver were assayed for fertility: **b** number of eggs laid per female in 24 hours; **c** fraction of viable eggs per female after 48h and **d** estimated progeny, given by the number of eggs multiplied by the fraction of viable eggs. Each point represents the Cohen’s *d* value for each genetic manipulation. Values below -0.8 and above 0.8 were classified as a large effect size and are highlighted in green. Values between -0.8 and 0.8 are represented in grey.

We next tested our hypothesis by assessing nutrient appetite in females with germline perturbations. Prior to the behavioural assays, females were maintained in a fully fed and mated state. We took advantage of the holidic medium (HM), a complete chemically defined diet to maintain the animals in a stable and consistent diet^8,17^. Feeding behavior was quantified using the fly Proboscis and Activity Detector (flyPAD), which provides automated, high-resolution measurements of feeding by measuring the number of proboscis–food contact events^35^ (“sips”, **Fig. 2a**). This screen identified 18 germline-specific gene knockdowns with measurable increases in yeast appetite, of which 15 were statistically significant (**Fig. 2b** and **d**, and **Supplementary Fig. 2a**). Only three additional knockdowns led to a significant decrease in yeast appetite (**Fig. 2b** and **d**, and **Supplementary Fig. 2a**). Furthermore, changes in yeast appetite did not correlate with changes in sugar appetite across manipulations (**Fig. 2b-e**, and **Supplementary Fig. 2a-b**). Although several manipulations also induced detectable changes in sugar appetite, these effects were weaker and less consistent than those observed for yeast (**Fig. 2c** and **Supplementary Fig. 2b**).

**Figure 2.**
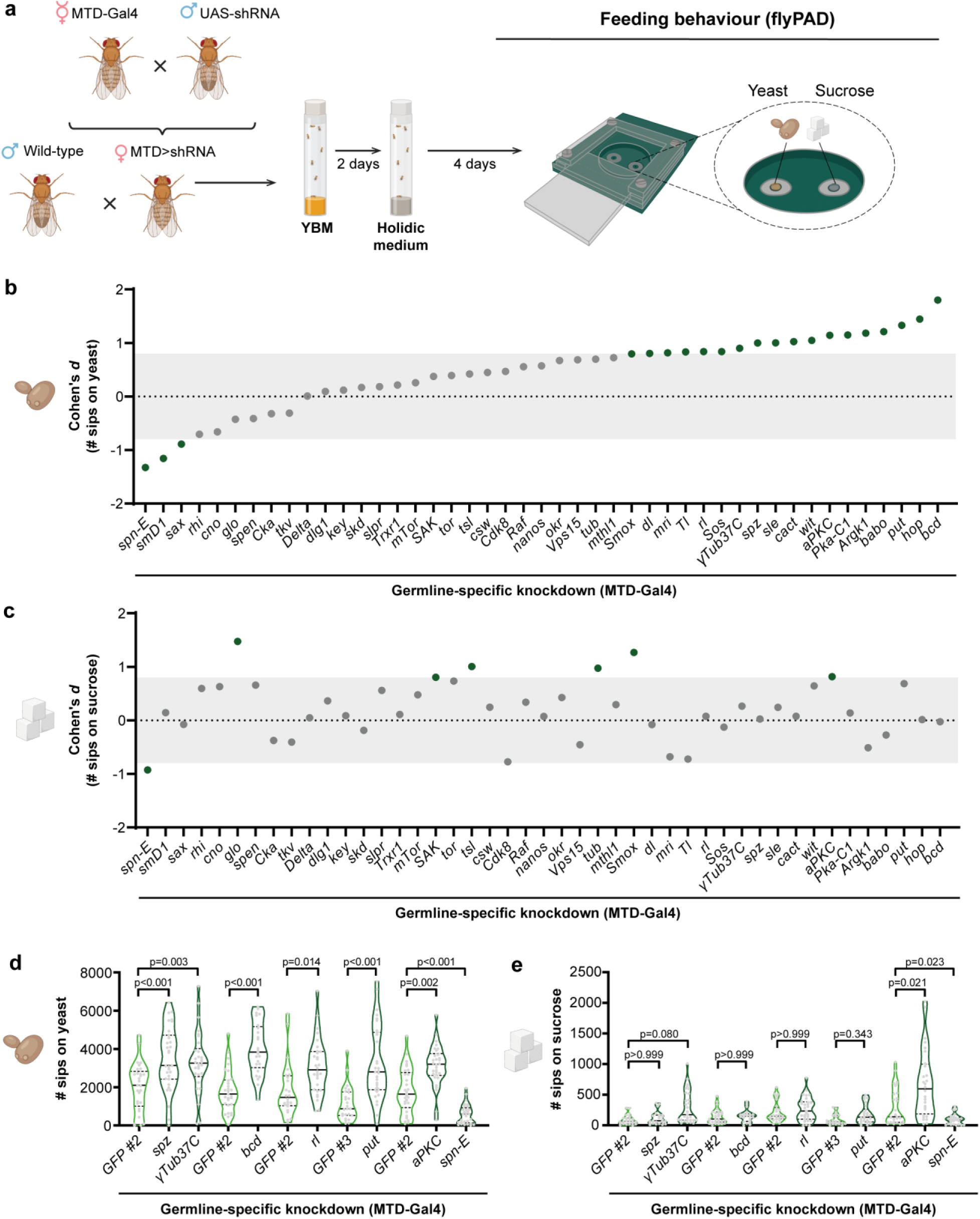
Ovarian physiology impacts yeast appetite. **a** Schematic representation of experimental and feeding behavioral set-ups. Further details can be found in the Methods section (YBM-Yeast Based Medium). **b**, **c** Fully fed mated females knocked down for the indicated genes using the MTD-Gal4 driver were assayed for feeding behavior using the FlyPAD: total number of sips on yeast (**b**) and sucrose (**c**). Each point represents the Cohen’s *d* value for each genetic manipulation. Values below -0.8 and above 0.8 were classified as a large effect size and are highlighted in green. Values between -0.8 and 0.8 are represented in grey. **d, e** Fully fed mated females knocked down for the indicated genes in the germline were assayed for feeding behavior using the FlyPAD. Total number of sips on yeast **(d)** or sucrose **(e)** are represented as truncated violin plots and each dot corresponds to one individual fly. Median and quartiles are represented in solid or dashed lines, respectively. *GFP* RNAi was used as negative control (GFP#2-valium 20; attP2; GFP#3 - valium 20; attp40). n = 24-31. The statistical significance was determined using ordinary one-way ANOVA or t-test in cases where the data followed a normal distribution, and the Kruskal-Wallis test or Mann-Whitney in the remaining cases.

Of the 17 candidate genes from the screen that produced a behavioral phenotype, we validated the effect for 10: *spz*, *aPKC*, *put*, *bcd*, *γTub37C*, *hop*, *dl*, *rl*, *sle* and *spn-E*. To further validate the strongest candidates, we repeated the gene knockdowns for 7 of these genes using *matα4*-Gal4, which drives expression in the germline only after stage 2–3 of egg chamber development, thereby sparing early oogenesis in the germarium. Using this driver, 6 out of the 7 knockdowns tested recapitulated the nutrient appetite phenotypes observed with MTD-Gal4 (**Supplementary Fig. 3**). These results indicate that the feeding phenotypes arise from disruptions in mid to late oogenesis rather than earlier germarial stages.

Yeast is a protein-rich food source for *Drosophila*, and increased yeast appetite is typically associated with elevated amino acid demands, either to restore homeostasis or to support reproduction upon mating^8,37,70,38–40,71,34,41^. To determine whether the feeding changes were specific to yeast or reflected a broader increase towards amino-acid rich foods, we measured total intake of a complete holidic medium using a dye-based ingestion assay (**Supplementary Fig. 4a**). Females with two different germline perturbations that increased yeast appetite also increased consumption of this complete diet supporting the idea that ovarian physiology regulates the appetite for nutritionally complete foods (**Supplementary Fig. 4b**).

Collectively, our results suggest the existence of an axis of communication originating in the ovaries that controls yeast appetite. Because most germline manipulations induced an increase in yeast appetite, we next focused on further characterizing this phenotype.

### Ovarian control of yeast appetite is independent of amino acid state but require mating

Previous work has begun to elucidate how yeast appetite is regulated in *Drosophila*, particularly in females. In these animals, protein deprivation, and more specifically, deprivation of essential amino acids, is a strong driver of yeast consumption^8,34,37–41,70,71^. Dietary protein and amino acids are also required to sustain female fertility^8,17,34,41–45^. Given this critical link between amino acid availability, reproductive physiology, and nutrient-specific appetite, we asked whether the increased yeast appetite observed upon germline perturbation depends on the animal’s amino acid state.

To test this, we selected a subset of germline-specific knockdowns that reliably increased yeast appetite and assayed their feeding behavior under amino acid-deprived conditions using the flyPAD. Surprisingly, similarly to what was observed in fully fed animals, germline-manipulated females deprived of amino acids also displayed an increase in yeast appetite when compared to control animals (**Fig. 2d**, **Fig. 3a**, **Supplementary Fig. 2a** and **Supplementary Fig. 5a**). For all manipulations tested, yeast consumption increased across diets indicating that the germline-dependent mechanism enhancing yeast appetite operates independently of amino acid deprivation pathways. The latter are likely systemic rather than ovary-derived and these regulatory mechanisms are thus additive.

**Figure 3.**
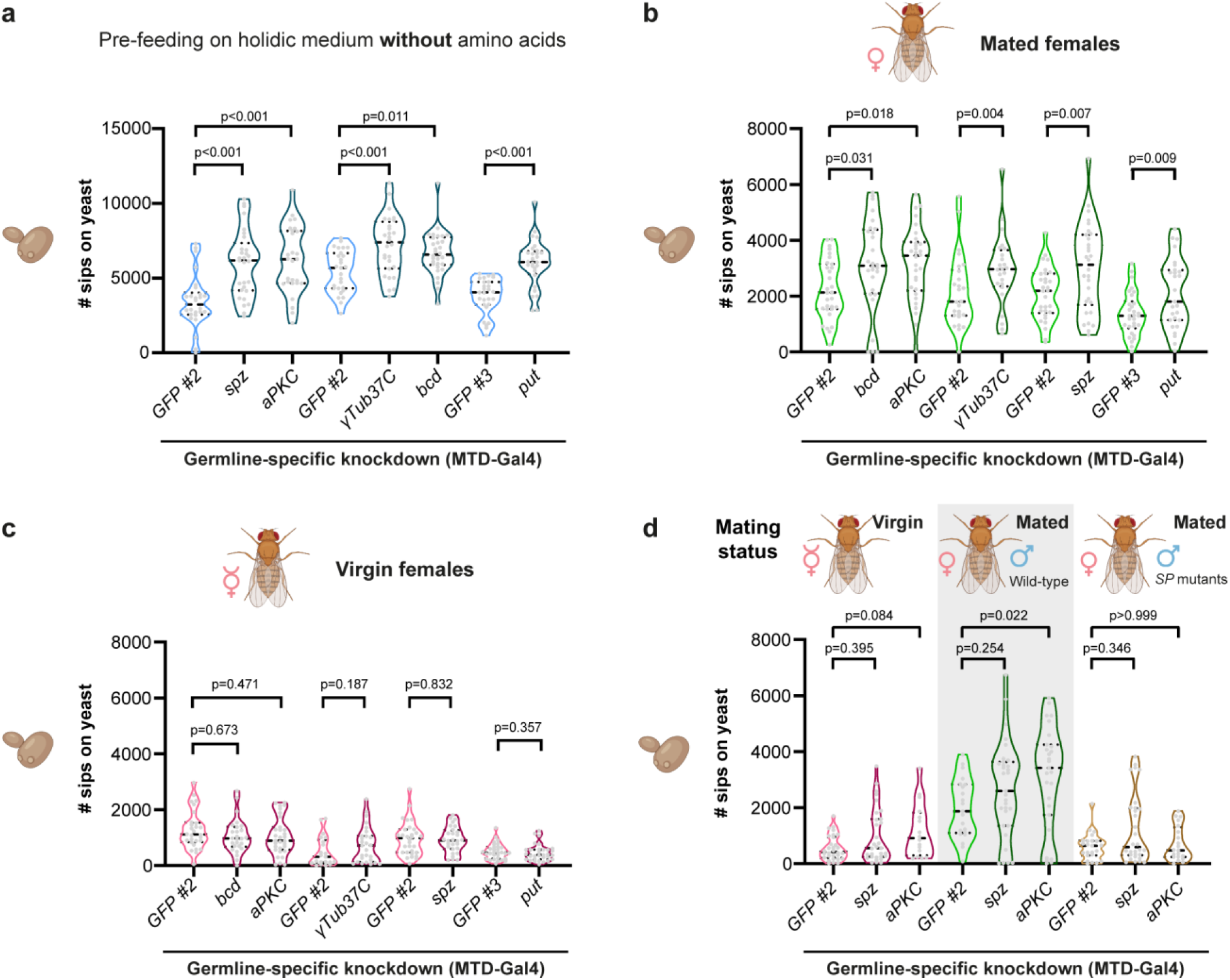
Ovarian control of yeast appetite is independent of amino acid state but requires mating. **a** Females knocked down for the indicated genes in the germline using the MTD-Gal4 driver were pre-treated for 4 days with a synthetic diet from where all amino acids were omitted. **b-d** Fully fed females mated with wild type or Sex Peptide (SP) mutant males **(b, d)** or virgins **(c, d)** knocked down for the indicated genes in the germline using the MTD-Gal4 driver were assayed for feeding behavior using the flyPAD**. a-d** Total number of sips on yeast are represented as truncated violin plots and each dot corresponds to one individual fly. Median and quartiles are represented in solid or dashed lines, respectively. *GFP* RNAi was used as negative control (GFP#2 - valium 20; attP2; GFP#3 - valium 20; attp40). n = 21-34. The statistical significance was determined using ordinary one-way ANOVA or t-test in cases where the data followed a normal distribution, and the Kruskal-Wallis test or Mann-Whitney in the remaining cases.

Mating state is another strong modulator of yeast appetite in *Drosophila* females. Mating increases yeast appetite, which is further strongly increased under conditions of protein deprivation^37–41^. Mating also triggers broad physiological alterations that enhance reproductive output, including increased egg production and ovulation^41,72–77^. Given this tight association between mating status, reproductive physiology, and yeast-specific appetite, we next asked whether the increase in yeast feeding observed in germline-manipulated females depended on their mating status.

Using the same subset of germline knockdowns, we collected virgin females and either maintained them isolated (virgin condition) or housed with males for six days to ensure a robust mated physiological state (mated condition). Feeding behaviour was then assessed using the flyPAD. Strikingly, germline-manipulated females displayed increased yeast appetite only when mated (**Fig. 3b-c** and **Supplementary Fig. 5b-c**). Virgin females carrying the same germline knockdowns showed no significant changes in yeast intake relative to controls (**Fig. 3c** and **Supplementary Fig. 5c**). These results show that the feeding phenotype of germline-manipulated females is dependent on the mating status.

During copulation, males transfer not only sperm but also an array of peptides and metabolites into the female reproductive tract^41,78–81^. These factors elicit widespread post-mating changes, many of which depend on sex peptide (SP), which binds to its receptor on Sex Peptide Receptor Neurons (SPSNs) located around the oviducts^41,72–77,82,83^. SP binding suppresses SPSN activity, triggering a neural cascade to the abdominal ganglion and subsequently the central brain, where it modulates diverse aspects of female physiology and behaviour, including nutrient-specific feeding^40,76,77,83–86^. Because increased yeast appetite in germline-manipulated females is only visible upon mating, we tested whether this effect requires SP signalling. We measured yeast appetite in females mated either with wild type or SP mutant males. While germline-manipulated females mated with wild-type males showed an increase in yeast appetite, this phenotype was largely abolished in females mated with SP mutants, mirroring the phenotype of virgin females (**Figure 3d** and **Supplementary Fig. 5d**).

Taken together, these results demonstrate that while the oogenesis-dependent modulation of yeast appetite is independent of the amino acid state, it critically requires the post-mating physiological state mediated by SP signalling.

### Ovarian and mating signals both increase yeast appetite by boosting feeding frequency

The fine structure of feeding behaviour has been used across species to reveal the behavioural mechanisms underlying homeostatic regulation^35,87–89^. High-resolution, time-resolved behavioural data can help separate specific components of feeding control, such as the drive to initiate versus terminate a meal^89,90^. Similarly, in *Drosophila*, individual sips are organized into feeding bursts that cluster together to form an activity bout^35^ (**Supplementary Fig. 6a**).

Previous studies have shown that both nutritional and mating states can alter distinct components of feeding microstructure during yeast consumption^35,39,40,91,92^. We reasoned that analyzing the microstructure of feeding in our conditions could reveal whether the ovarian mechanism we uncovered acts through known behavioral pathways or engages a distinct one. To identify which parameter best explained the changes in yeast appetite observed in germline-manipulated females, we performed correlation analyses between individual microstructure parameters and the total number of sips on yeast. The number of feeding bursts showed a strong positive correlation with total yeast intake (**Supplementary Fig. 6b-c**). Coherent with the idea that germline perturbed animals initiate feeding more often resulting in higher number of feeding bursts, we also find a consistent reduction of inter-burst interval length (IBI’s) in the germline manipulated animals when compared to the control (**Supplementary Fig. 6d-e**). In contrast, other parameters including burst duration or number of sips per burst, did not show robust correlations with total number of sips (**Supplementary Fig. 6f-i**).

To understand how this pattern compares with canonical drivers of yeast appetite, we analysed microstructural changes induced by amino-acid deprivation and mating. As reported in the literature, amino acid–deprived females showed broad alterations in feeding microstructure: increased sips per burst, resulting in longer bursts overall, as well as reduced IBIs, which together produced more total feeding bursts^39,40,92^ (**Supplementary Fig. 7a**). Thus, amino-acid deprivation acts through multiple adjustments to both burst structure and burst frequency. In contrast, the increase in yeast appetite in fully fed mated females was explained primarily by an increase in the number of feeding bursts accompanied by shorter IBIs, with less pronounced changes in burst duration or sip number per burst (**Supplementary Fig. 7b**).

Interestingly, germline-manipulated females did not consistently exhibit increases in sips per burst or burst duration relative to controls. Instead, their elevated yeast appetite was primarily associated with shortened IBIs and a corresponding increase in the number of feeding bursts, reflecting key aspects of the post-mating feeding profile (**Supplementary Fig. 6** and **7b**).

Together, these results indicate that ovarian regulation of yeast appetite relies predominantly on mechanisms that increase the initiation of feeding bursts. This strategy is at least partially distinct from the one engaged by amino-acid deprivation, which alters multiple aspects of feeding microstructure. The parallels with the post-mating feeding phenotype support the idea that ovarian signals, and mating-derived cues, converge on shared or interacting mechanisms that modulate nutrient-specific feeding behaviour.

### Changes in yeast appetite reflect delays in oogenesis progression

Our behavioral analyses of females with germline-specific perturbations establish a correlation between ovarian physiology and the regulation of nutrient-specific feeding. This raised a key question: which ovarian states and ovarian signal(s) drives the changes in yeast appetite that we observe?

Despite the diverse functions of the genes whose knockdown consistently increased yeast appetite, including transcription factors, kinases, cytoskeletal elements, and germline regulators, their biological roles converge on processes essential for oocyte and embryonic patterning and signaling^93–106^. Many are maternally expressed and are required during oogenesis or early embryonic development^93–106^. This convergence led us to hypothesize that the altered feeding phenotypes may arise from shared defects in oogenesis caused by these manipulations.

To first determine whether global fertility defects alone could explain the changes in yeast appetite, we established a correlation between the median number of sips on yeast with three fertility measures: median egg number, egg viability, and estimated progeny for each candidate gene from the screen (**Fig. 4a-c**). We detected a negative correlation between both yeast intake and egg production; and yeast intake and egg viability, reaching statistical significance with the latter (**Fig. 4a-c, Supplementary Fig. 1** and **Supplementary Fig 2**). However, the relatively low r² values indicate that fertility defects explain only a portion of the variation in feeding behavior. Importantly, several manipulations altered fertility without affecting yeast appetite, and vice versa, suggesting that there was no strict functional link between fertility per se and yeast appetite (**Fig. 1**, **Fig.2**, **Fig. 4a-c, Supplementary Fig. 1** and **Supplementary Fig 2**).

**Figure 4.**
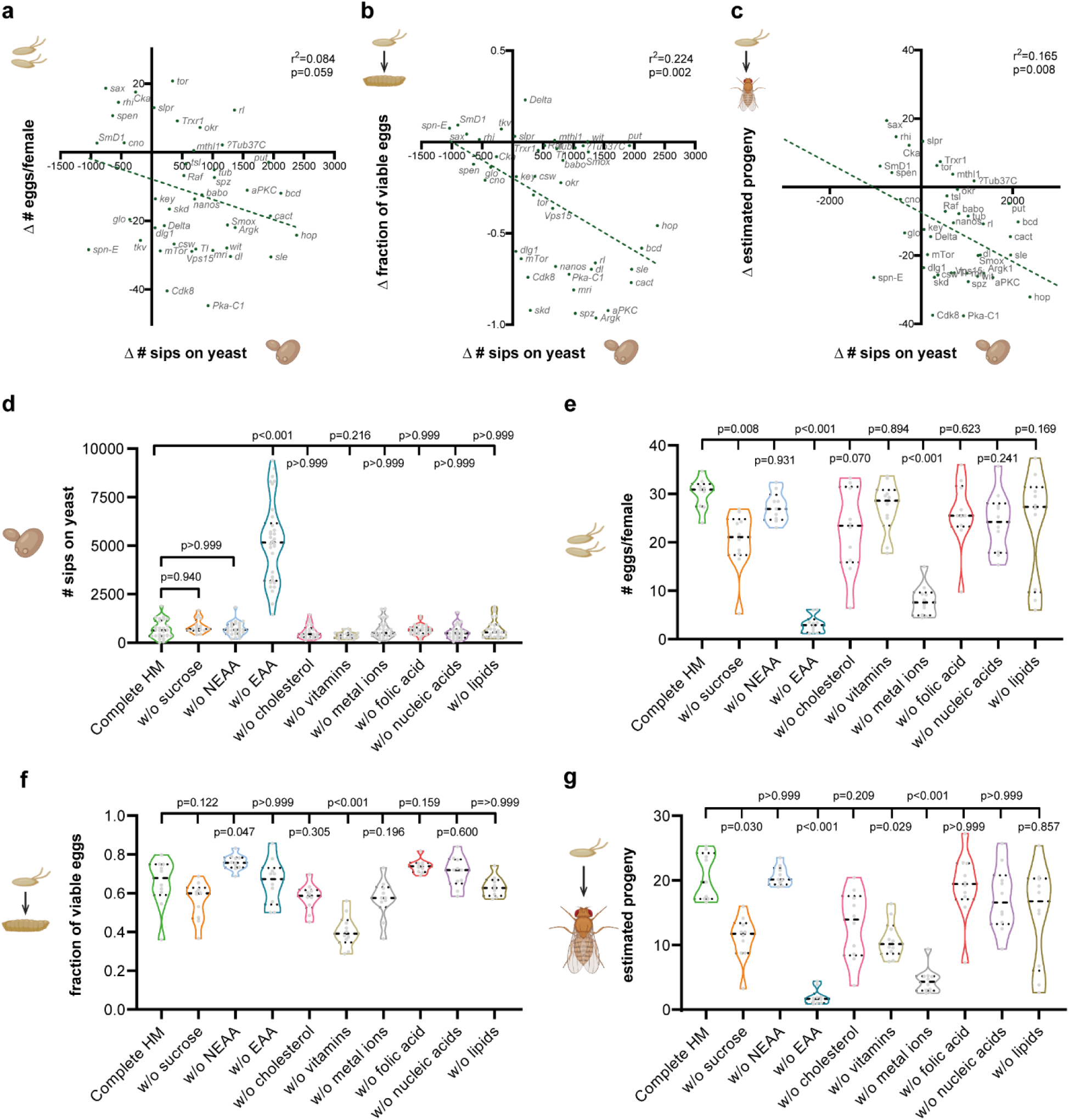
Reduced female fertility is not enough to drive yeast appetite. **a-c** Correlation between the number of sips on yeast and the number of eggs laid per female (**a**), the fraction of viable eggs (**b**) or the estimated progeny (**c**). Each point represents a germline specific gene-knockdown. The regression line represents the best fit for the data. For further details, see Materials and Methods. **d-g** Wild-type mated females were pre-treated for 4 days with either a complete holidic medium, or a holidic medium from where specific groups of nutrients were omitted (w/o – without; EAA - essential amino acids; NEAA - non-essential amino acids). Complete holidic medium (HM) is used as negative control. Median and quartiles are represented in solid or dashed lines, respectively. **d** Females were assayed for feeding behavior using the flyPAD. Total number of sips on yeast are represented as truncated violin plots and each dot corresponds to one individual fly. n = 16-30. **e-g** Females were assayed for fertility, by calculating the number of eggs laid per female (**e**), the fraction of viable eggs (**f**), and the estimated progeny (**g**). **d-g** Data are represented as truncated violin plots and median and quartiles are represented in solid or dashed lines, respectively. n = 9-10. The statistical significance was determined using ordinary one-way ANOVA in cases where the data followed a normal distribution and the Kruskal-Wallis test in the remaining cases.

To further test whether reduced progeny output could lead to an increase in yeast appetite, we tested whether other manipulations known to affect egg production also led to a similar phenotype. Because multiple dietary components influence reproduction, we asked whether deprivation of these nutrients similarly modulates yeast appetite. We therefore deprived wild-type females of specific nutrient groups for four days and assayed feeding behavior using the flyPAD. We also measured fertility, by monitoring egg number and viability (**Fig. 1a and 2a**).

In line with previous work using two-color preference assays, removal of essential amino acids was the only nutritional manipulation that specifically increased yeast appetite^34^ (**Fig. 4d** and **Supplementary Fig. 8**). In contrast, reduced progeny output, whether due to a decline in egg number or viability, was observed across a broad range of nutrient deprivations, not only essential amino acids; including sucrose, vitamins and metal ions^8^ (**Fig. 4d-g**). Thus, although several dietary nutrients are required for optimal egg production, essential amino acid deprivation uniquely impacts both reproduction and yeast appetite. These results further corroborate that while reproductive output and yeast appetite are related, feeding changes in germline-manipulated females are unlikely to be explained simply by a reduction in progeny production.

This led us to consider whether specific defects in oogenesis progression, rather than global fertility reduction, might underlie the observed feeding phenotypes. To test this, we examined ovarian morphology in females with germline-specific knockdown of two genes (*aPKC* and *spz*) that produced robust increases in yeast appetite. Ovaries were stained with phalloidin to visualize the actin architecture, allowing staging and quantification of individual egg chambers (**Fig. 5a** and **Supplementary Fig. 9**).

**Figure 5.**
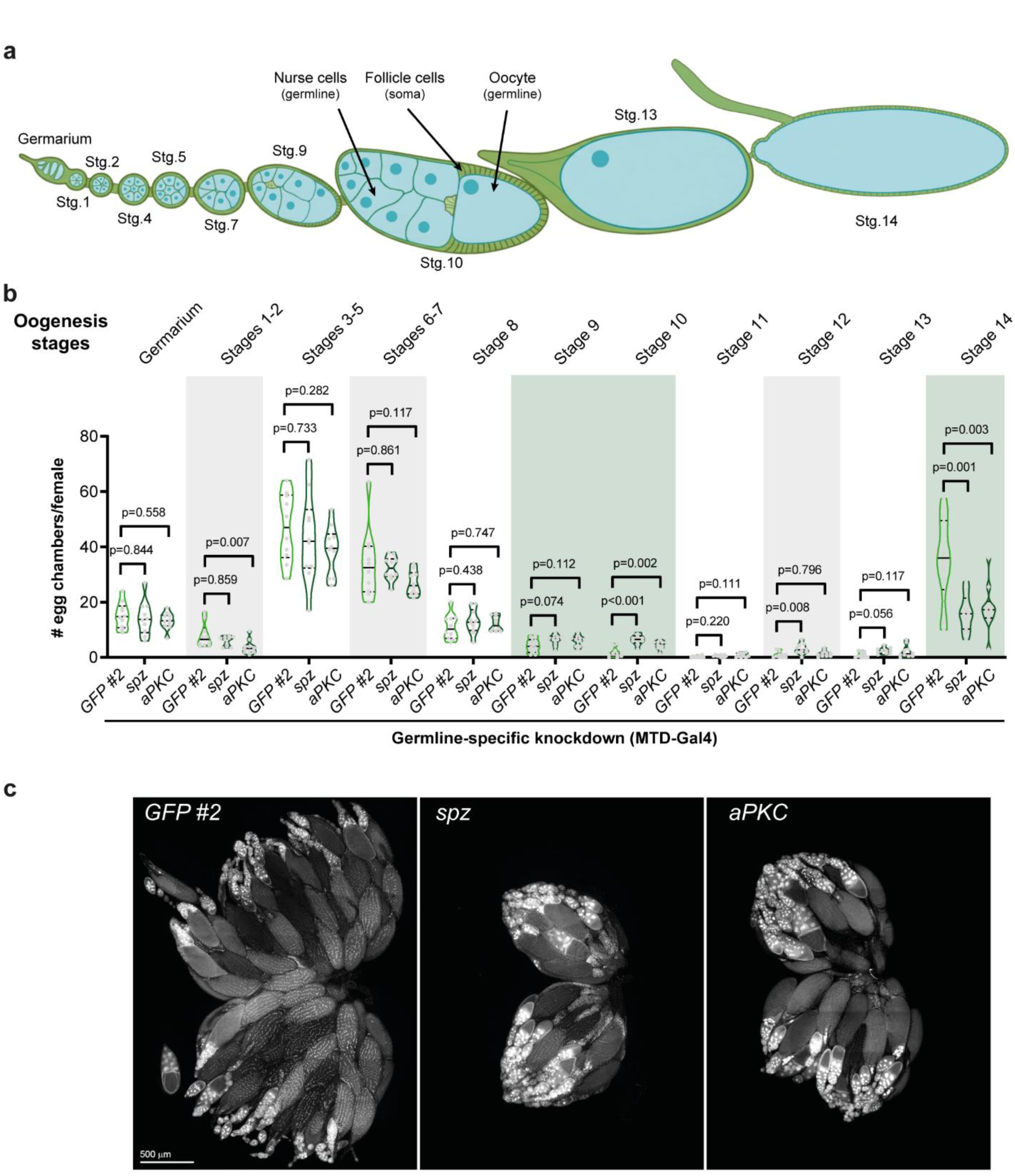
Germline perturbations that affect yeast appetite commonly alter oogenesis progression. **a** Schematic representation of the egg chamber development within the ovariole. Each ovariole consists of various egg chambers at different developmental stages (Stg). **b** Fully fed mated females knocked down for the indicated genes using the MTD-Gal4 driver were assayed for the number of egg chambers in each phase of oogenesis. Total number of egg chambers per stage per female are represented as truncated violin plots and median and quartiles are represented in solid or dashed lines, respectively. *GFP* RNAi was used as negative control (GFP#2 - valium 20; attP2). n =10. The statistical significance was determined using ordinary one-way ANOVA in cases where the data followed a normal distribution, and the Kruskal-Wallis test in the remaining cases. **c** Ovary morphology revealed by immunostaining of ovaries from females where *spz* or *aPKC* were knocked down in the germline using the MTD-Gal4 driver. Ovaries were stained with DAPI (DNA). Scale bar, 500 μm.

Strikingly, knockdown of either *aPKC* or *spz* in the germline caused similar disruptions in oogenesis progression (**Fig. 5b**). Specifically, both manipulations resulted in: i) an increased number of stage 9–10 egg chambers (significant at stage 10); and ii) a pronounced reduction in the number of mature stage 14 oocytes (**Fig. 5b-c**). To determine whether this phenotype was generalizable, we extended the analysis to 2 additional germline knockdowns that increased yeast appetite. These germline manipulations showed a similar pattern: an accumulation of stage 10 egg chambers and a consistent decrease in stage 14 oocytes (**Supplementary Fig. 10**).

Together, these findings strongly suggest that germline perturbations that delay or partially arrest oogenesis at stages 9-10 while decreasing the number of stage 14 oocytes, lead to elevated yeast appetite. This shared physiological signature suggests that the animal monitors the progression of oogenesis and adjusts nutrient-specific appetite accordingly, allowing for the coordination of feeding behavior with reproductive state.

### Ovary-derived Dilp8 signaling regulates yeast appetite in mated females

Because stage-14 oocytes were consistently reduced in females with germline knockdown of *aPKC* or *spz*, we investigated whether signaling molecules normally expressed at this late stage might also be reduced in these ovaries. Follicle cells of stage-13–14 egg chambers are known to express *Drosophila* insulin-like peptide 8 (Dilp8), a relaxin-like peptide^107–115^. Consistent with a reduction in mature oocytes, ovaries from *aPKC* and *spz* germline knockdowns showed significantly decreased *dilp8* transcript levels (**Fig. 6a**). This reduction suggests that Dilp8 may function as an ovarian inhibitory signal that restrains yeast appetite.

**Figure 6.**
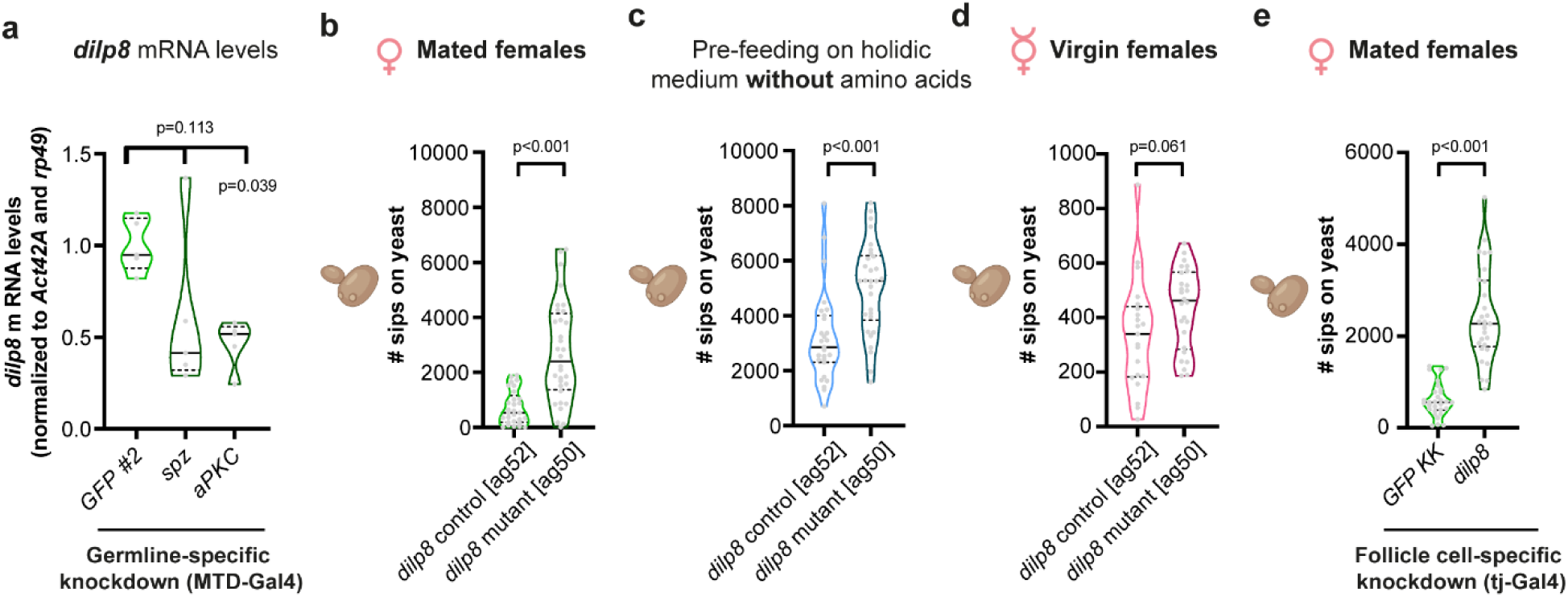
*dilp8* expression from ovarian follicle cells regulates yeast appetite. **a** Ovaries from fully fed mated females knocked down for the indicated genes using the MTD-Gal4 driver were dissected and RNA extracted for assessing *dilp8* expression levels. *GFP* RNAi was used as negative control (GFP#2 - valium 20; attP2). Normalized *dilp8* mRNA levels are represented as truncated violin plots with each dot corresponding to an independent sample generated from 8 ovaries and median and quartiles are represented in solid or dashed lines, respectively. n = 5. The statistical significance was determined using the Kruskal-Wallis test. *dilp8* control or knockout mated fully fed **(b)**, mated amino-acid deprived **(c)**, or fully fed virgins **(d)**; or females knocked down for the *dilp8* in the follicle cells **(e)** were assayed for feeding behavior using the flyPAD. **b-e** Total number of sips on yeast are represented as truncated violin plots and each dot corresponds to one individual fly. Median and quartiles are represented in solid or dashed lines, respectively. The corresponding genetic background control was used as control for *dilp8* knockout mutant (**b-d**) and *GFP* RNAi was used as negative control (**e**). n = 21-32. The statistical significance was determined using t-test in cases where the data followed a normal distribution and Mann-Whitney in the remaining cases.

To test this hypothesis directly, we examined the feeding behavior of *dilp8* mutant females. As predicted, mated *dilp8* mutants exhibited a robust and significant increase in yeast appetite, resembling the phenotype observed in the germline-manipulated females (**Fig. 6b** and **Supplementary Fig. 11a**). This phenotype was also independent of internal amino acid state, as females deprived from amino acids showed the expected increase in yeast appetite, and there was an additive effect of *dilp8* mutation (**Fig. 6c** and **Supplementary Fig. 11b**). Interestingly, and consistent with the behavior of germline-manipulated females, the elevated yeast appetite of *dilp8* mutants was strictly mating-dependent (**Fig. 6b** and **d**, and **Supplementary Fig. 11a** and **c**). To determine whether this mechanism requires specifically the pool of ovarian Dilp8, we knocked down *dilp8* in follicle cells. These females also showed increased yeast intake supporting the idea that follicle cell–derived Dilp8 mediates the ovarian regulation of yeast appetite (**Fig. 6e** and **Supplementary Fig. 11d**).

Taken together, these results support a model in which yeast appetite is tuned by the abundance of late-stage oocytes through a Dilp8-dependent signaling pathway that is mechanistically distinct from amino acid–driven homeostatic feeding. In this framework, accumulation of mature (stage-14) oocytes promotes increased Dilp8 secretion from follicle cells, which in turn suppresses yeast appetite. Conversely, when oogenesis is delayed and stage-14 oocytes are reduced, Dilp8 levels fall, leading to elevated yeast consumption, particularly in mated females.

## Discussion

Regulation of food intake is essential for ensuring that animals obtain the nutrients required to sustain physiological functions while avoiding harmful overconsumption. Although decades of research have identified many molecules and neural pathways involved in feeding control, how individual organs communicate their physiological state to regulate nutrient-specific appetites remains incompletely understood. Most studies have focused on direct detection of missing nutrients, such as amino acid or sugar. Far less is known about how the physiological status of organs shape appetites in the absence of evident nutritional deficiency. Here, we identify a previously unrecognized communication axis originating in the ovary that modulates yeast feeding, revealing a new mechanism through which organ-specific demands tune nutrient-specific appetite (**Fig. 7**).

**Figure 7.**
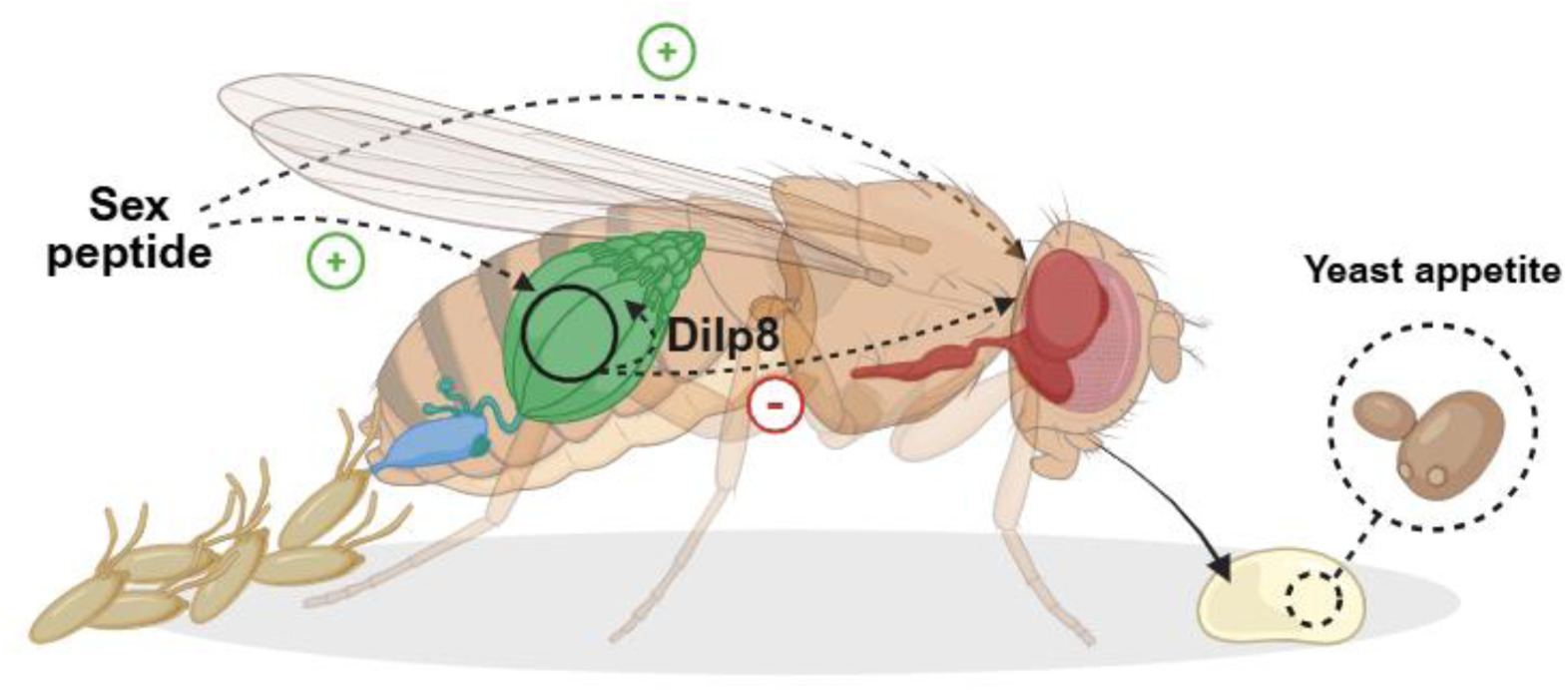
Ovarian control of yeast-specific appetite. Sex Peptide transferred during mating stimulates oogenesis, ovulation, and activates the mating circuit, leading to increased yeast appetite. The ovarian–feeding axis operates specifically in mated females and involves the inhibitory action of Dilp8 produced by late stage follicles. Dilp8 may regulate yeast appetite through two non-mutually exclusive mechanisms: (a) by locally restraining oogenesis, thereby limiting the accumulation of nutrient demanding pre vitellogenic and vitellogenic follicles thereby reducing a positive, appetite promoting signal; or b) by acting directly on the central nervous system to dampen activity in the mating dependent feeding circuit. Green “+”, positive effect; red “-”, inhibitory effect; dashed arrows, inhibitory effect; dashed arrows indicate multi step or indirect communication pathways. Created in BioRender. Santos, Z. (2026) https://BioRender.com/ss0uqqy.

Our findings demonstrate that yeast-specific appetite is regulated by the physiological state of the ovary and depends on oogenesis progression. This regulation is engaged only in reproductively active (mated) females. Germline perturbations that alter oocyte development consistently shift yeast intake, indicating that the ovary conveys information about its functional state to central feeding circuits. How this communication is achieved remains unclear. Below, we outline two potential models for how this regulatory axis might function.

### Regulation of yeast-specific appetite by the ovary

Our initial results show that several germline manipulations lead to an increased yeast appetite. According to the literature, such manipulations, including *aPKC*, *spz*, *bcd*, and *γtub37C* RNAi, are predicted to disrupt processes essential for early- and mid-oogenesis (e.g., oocyte specification, polarity establishment, or RNA-based regulation). These processes are critical for proper oogenesis progression, likely explaining the accumulation of mid-stage follicles and reduction of late-stage follicles observed upon gene knockdown^93–106^. Because RNAi typically produces hypomorphic rather than null phenotypes, these perturbations may only partially disrupt oogenesis allowing it to proceed, albeit with altered dynamics^116–118^. This nuance may have been essential for uncovering the communication mechanism we describe, as stronger disruptions would likely cause complete ovarian arrest masking the signaling logic.

A decrease in late-stage oocytes in these manipulations pointed us toward Dilp8, a relaxin-like peptide secreted by mature follicle cells. We showed that these manipulations led to a reduction in ovarian Dilp8 expression, consistent with their reduced number of mature oocytes. Furthermore, knockout or follicle cell-specific knockdown of Dilp8 phenocopies the feeding phenotypes of the germline knockdowns. In this context, Dilp8 therefore emerges as a key component of an inhibitory arm of the ovarian-feeding axis, acting as a yeast-specific satiety signal, keeping yeast intake within an optimal range.

Importantly, this inhibitory arm operates only in mated females. Virgin females, as well as females mated to SP mutant males, lack the increase in yeast appetite normally elicited by ovarian perturbations. SP induces long-lasting post-mating responses in females, including increased oogenesis, ovulation, and behavioral changes including an anticipatory increase in yeast appetite^37,40,76,83–86^. In doing so, SP functions as the primary driver of yeast appetite in females, directly acting at the central nervous system. But additionally, it also licenses the ovarian signal by: a) stimulating oogenesis, leading to an increase in the number of pre-vitellogenic and vitellogenic egg chambers, thereby increasing the nutritional requirements of the ovary; and b) by increasing ovulation, SP reduces the pool of Dilp8-expressing follicles, releasing the Dilp8-dependent satiety signal. This logic is intuitive: virgin females have low ovarian activity and therefore little need to adjust protein appetite, whereas mated females must align nutrient intake with rapidly increasing reproductive demands. SP provides an anticipatory signal, independent of ovarian physiology, while ovarian derived signals fine tune feeding according to current ovarian needs and restraining the female from overeating.

How might ovarian signals modulate yeast appetite? Beyond its roles in larval growth coordination and pupariation, recent work shows that Dilp8 also prevents excessive accumulation of mature follicles, maintaining oocyte quality in virgin females^107–115^. Mechanistically, Dilp8 is proposed to restrain oogenesis progression, limiting the number of follicles entering vitellogenesis, and promoting ovulation through its receptor, Lgr3, in neurons innervating the oviducts, thereby counteracting the ovarian SP functions^114,115^. In the context of feeding, Dilp8 could therefore act as a buffering signal: by restraining entry into vitellogenesis, it indirectly reduces the ovarian nutritional demands. When Dilp8 levels decrease, this restraint is lost, generating a state of enhanced protein demand.

Another possibility is that Dilp8 acts systemically. The function of specific neuronal populations expressing Lgr3 has been characterized in central brain. Lgr3^+^ neurons in the superior medial protocerebrum, forming part of the mating-dependent feeding circuit, have been shown to promote sugar intake in mated females^119^. It is thus possible that Dilp8 acts on these, or on other yet-unidentified neuronal populations, to influence appetite for protein-rich foods.

But how are ovarian needs conveyed to the nervous system? Multiple lines of evidence indicate that there is likely a positive ovarian signal regulating feeding, which can act independently, or be modulated by Dilp8 local ovarian functions. We have previously shown that complete germline ablation or severe disruption of mid-to-late oogenesis, both expected to drastically reduce or fully ablate Dilp8 levels since late stage oocytes are not produced, do not lead to an increase in yeast appetite^36,40,120^. This suggests the existence of an additional appetitive ovarian signal that promotes yeast feeding. Our data suggests that early vitellogenic follicles (stages 8–10) may produce such a positive cue, reflecting their substantial biosynthetic demand. One compelling candidate for such positive ovarian signal is ecdysone, a steroid hormone synthesized by follicle cells and known to coordinate multiple aspects of female reproduction^41,72,121–125^. Previous studies have implicated ecdysone signaling in modulating feeding behavior by acting directly in the central nervous system, consistent with the idea that early vitellogenic follicles could promote yeast appetite through steroid production^50^.

Independently of which and how the signal(s) reaches the central nervous system, our feeding microstructure analysis show that such signal(s) primarily affect the initiation of yeast feeding, a pattern that mirrors mating-induced changes and differs from amino acid deprivation responses. This strongly indicates that Dilp8 directly and/or other unidentified ovarian-originated signal(s) are likely to act at the level of the SAG mating circuit, either by gating its activity, or acting downstream.

### Ecological and physiological relevance of the ovarian feeding axis

Yeast is a nutritionally rich food source that provides amino acids, sterols, vitamins, and micronutrients essential for oogenesis. In conditions where oogenesis progression is delayed and vitellogenic follicles accumulate, an increase in yeast appetite may simply reflect the increased nutritional demand required to support continued follicle growth. This demand is met cooperatively with the fat body, the *Drosophila* counterpart of adipose tissue, which supplies yolk precursors and other metabolic substrates during vitellogenesis.

In natural environments, flies can encounter foods that are not calorically limiting but are suboptimal for ovarian physiology, causing localized stress or developmental delays without inducing systemic starvation. The ovary is particularly sensitive to environmental quality: plant toxins, microbial pathogens, and xenobiotics can disrupt oogenesis progression and/or ovulation even when overall nutrition is adequate^126–136^. Subtle disruptions in oogenesis derived from non-optimal food substrates could therefore elicit compensatory feeding behaviors.

From this eco-physiological perspective, our findings suggest that the ovary functions as a sensitive internal sensor that detects oogenesis perturbations triggering increased yeast consumption to promote reproductive recovery and enhance the animal’s resilience to environmental stressors. The ovary thus operates not only as a reproductive tissue but also as an integrator of physiological and environmental information, translating perturbations in oogenesis into adaptive behavior responses.

### Broader implications

Together, our findings uncover a previously unrecognized mechanism by which the ovary shapes yeast-specific appetite, establishing oogenesis progression rather than fertility or systemic nutrient deficiency, as a key internal variable that adjusts yeast feeding in a mating-dependent manner. Although demonstrated in *Drosophila*, this conceptual framework is likely to extend beyond insects. In mammals, ovarian physiology is known to influence food intake across the estrous and menstrual cycles, yet the nutrient specificity and underlying molecular logic remain poorly defined^137–145^. Our findings offer a mechanistic entry point for understanding how ovarian state influences discrete components of feeding behavior.

More broadly, the well-established association between ovarian dysfunction, metabolic disease, and obesity raises the possibility that mis regulation of ovarian signals could directly contribute to metabolic imbalance rather than merely resulting from it. By revealing an ovarian communication axis that couples reproductive state to nutrient selection, this study provides a fresh perspective on the reciprocal relationship between reproduction and metabolism and suggests that subtle alterations in ovarian physiology may have far-reaching consequences for feeding behavior and metabolic health. Pinpointing the molecular identity of the ovarian signals involved and defining how they intersect with conserved mammalian circuits will be an important direction for future research.

## Acknowledgements

We thank the members of the Organismal Metabolic Physiology Laboratory, Silvia Henriques and other members of the Behavior and Metabolism Laboratory for helpful discussions and comments on the manuscript. We thank Célia Baltazar, Inês de Haan Vicente and Ana Paula Elias for technical support. We thank the technical support of the Bioimaging Platform of the Gulbenkian Institute for Molecular Medicine, funded by PPBI-POCI-01-0145-FEDER-022122. We also acknowledge the technical support of the Fly Platform (LISBOA-01-0145-FEDER-022170), Glass Wash and Media Preparation, and Hardware and Software Platform at Champalimaud Foundation. Stocks obtained from the Bloomington Drosophila Stock Center (NIH P40OD018537) were used in this study. Stocks from the TRiP RNAi collection were used in this study (NIGMS P41 GM132087: “Functional genomics resources for the Drosophila and broader research communities”; Office of the Director R24 OD030002: “TRiP resources for modeling human disease”).

Research in Z.C.S lab is supported by funding from Gulbenkian Institute for Molecular Medicine (GIMM), from Portuguese Foundation for Science and Technology (FCT) CEECINST/00021/2021/CP1771/CT0002, from the Chan Zuckerberg Initiative (CZI) DAF, an advised fund of Silicon Valley Community Foundation (Grant agreement No. 2023-331947), and from the European Union (ERC grant agreement No. 101043068 – SweetEggs (views and opinions expressed are however those of the author(s) only and do not necessarily reflect those of the European Union or the European Research Council Executive Agency. Neither the European Union nor the granting authority can be held responsible for them).

This project was funded in part by grant number GCRLE-1420 from the Global Consortium for Reproductive Longevity and Equality through the Buck Institute, made possible by the Bia-Echo Foundation to C.R. and Z.C.S.. The project leading to these results has also received funding from “la Caixa” Banking Foundation to C.R. under the project codes HR-17-00539 and HR23-00516, from the Chan Zuckerberg Initiative (CZI) DAF, an advised fund of Silicon Valley Community Foundation (Grant agreement No. 2023-331943). This project was supported by the Portuguese Foundation for Science and Technology (FCT) postdoctoral fellowship SFRH/BPD/79325/2011 to Z.C-S. Work by Z.C-S. was also financed by national funds through the FCT, in the framework of the financing of the Norma Transitória (DL 57/2016). Research at the Centre for the Unknown is supported by the Champalimaud Foundation.

Research in the A.G. lab was funded by the FCT (2022.03859.PTDC; 2023.15344.PEX), by the Research and Development Units iNOVA4Health (10.54499/UIDB/04462/2020) and Centre for Ecology, Evolution and Environmental Changes (cE3c) (10.54499/UIDB/00329/2020; 10.54499/UIDB/00329/2025), and by LS4FUTURE and CHANGE financed by the FCT/Ministério da Ciência, Tecnologia e Ensino Superior (Portugal).

## Notes

### Competing Interest Statement

The authors have declared no competing interest.

## References

1. Sun, F. & Poss, K. D. Inter-organ communication during tissue regeneration. Development 150, dev202166 (2023).

2. Droujinine, I. A. & Perrimon, N. Interorgan Communication Pathways in Physiology: Focus on *Drosophila*. Annu. Rev. Genet. 50, 539–570 (2016).

3. Tokizane, K. & Imai, S. Inter-organ communication is a critical machinery to regulate metabolism and aging. Trends in Endocrinology & Metabolism 36, 756–766 (2025).

4. Castillo-Armengol J., Fajas, L. & Lopez-Mejia, I. C. Inter-organ communication: a gatekeeper for metabolic health. EMBO Rep 20, (2019).

5. Priest, C. & Tontonoz, P. Inter-organ cross-talk in metabolic syndrome. Nat Metab 1, 1177–1188 (2019).

6. Münch, D., Ezra-Nevo, G., Francisco, A. P., Tastekin, I. & Ribeiro, C. Nutrient homeostasis — translating internal states to behavior. Current Opinion in Neurobiology 60, 67–75 (2020).

7. Lee, G., Lee, J., Suh, G. S. B. & Oh, Y. Post ingestive systemic nutrient sensing for whole-body homeostasis. Molecules and Cells 48, 100271 (2025).

8. Piper, M. D. W. et al. A holidic medium for Drosophila melanogaster. Nat Methods 11, 100–105 (2014).

9. Solon-Biet, S. M. et al. Macronutrient balance, reproductive function, and lifespan in aging mice. Proc Natl Acad Sci USA 112, 3481–3486 (2015).

10. Solon-Biet, S. M. et al. The Ratio of Macronutrients, Not Caloric Intake, Dictates Cardiometabolic Health, Aging, and Longevity in Ad Libitum-Fed Mice. Cell Metabolism 19, 418–430 (2014).

11. Solon-Biet, S. M. et al. Branched-chain amino acids impact health and lifespan indirectly via amino acid balance and appetite control. Nat Metab 1, 532–545 (2019).

12. Wali, J. A. et al. Determining the metabolic effects of dietary fat, sugars and fat-sugar interaction using nutritional geometry in a dietary challenge study with male mice. Nat Commun 14, 4409 (2023).

13. Di Francesco, A. et al. Dietary restriction impacts health and lifespan of genetically diverse mice. Nature 634, 684–692 (2024).

14. Morland, F. et al. Distinct macronutrient ratios optimize offspring survival, growth, and maternal glucose tolerance across mouse reproduction. Proc. Natl. Acad. Sci. U.S.A. 123, e2513044123 (2026).

15. Strilbytska, O. et al. Life-History Trade-Offs in Drosophila: Flies Select a Diet to Maximize Reproduction at the Expense of Lifespan. J Gerontol A Biol Sci Med Sci 79, glae057 (2024).

16. Lee, K. P. et al. Lifespan and reproduction in Drosophila: New insights from nutritional geometry. Proceedings of the National Academy of Sciences 105, 2498–2503 (2008).

17. Piper, M. D. et al. Matching Dietary Amino Acid Balance to the In Silico-Translated Exome Optimizes Growth and Reproduction without Cost to Lifespan. Cell Metab 25, 610–621 (2017).

18. Senior, A. M. et al. Multidimensional associations between nutrient intake and healthy ageing in humans. BMC Biol 20, 196 (2022).

19. Yap, Y. W. et al. Restriction of essential amino acids dictates the systemic metabolic response to dietary protein dilution. Nat Commun 11, 2894 (2020).

20. Brüning, J. C. & Fenselau, H. Integrative neurocircuits that control metabolism and food intake. Science 381, eabl7398 (2023).

21. Timper, K. & Brüning, J. C. Hypothalamic circuits regulating appetite and energy homeostasis: pathways to obesity. Disease Models & Mechanisms 10, 679–689 (2017).

22. Itskov, P. M. & Ribeiro, C. The Dilemmas of the Gourmet Fly: The Molecular and Neuronal Mechanisms of Feeding and Nutrient Decision Making in Drosophila. Front. Neurosci. 7, (2013).

23. Mahishi, D., Agrawal, N., Jiang, W. & Yapici, N. From Mammals to Insects: Exploring the Genetic and Neural Basis of Eating Behavior. Annual Review of Genetics 58, 455–485 (2024).

24. Nässel, D. R. & Zandawala, M. Recent advances in neuropeptide signaling in Drosophila, from genes to physiology and behavior. Progress in Neurobiology 179, 101607 (2019).

25. Nässel, D. R. & Zandawala, M. Hormonal axes in Drosophila: regulation of hormone release and multiplicity of actions. Cell Tissue Res 382, 233–266 (2020).

26. Yu, J. H. & Kim, M.-S. Molecular Mechanisms of Appetite Regulation. Diabetes Metab J 36, 391–398 (2012).

27. Simpson, S. J. & Abisgold, J. D. Compensation by locusts for changes in dietary nutrients: behavioural mechanisms. Physiological Entomology 10, 443–452 (1985).

28. Simpson, S. J., Couteur, D. G. L. & Raubenheimer, D. Putting the Balance Back in Diet. Cell 161, 18–23 (2015).

29. Simpson, S. J. & Raubenheimer, D. The Nature of Nutrition. (Princeton, Princeton University Press, 2012).

30. Gould, J. L. A Physiological Ethology: The Hungry Fly. A Physiological Study of the Behavior Associated with Feeding. V. G. Dethier. Harvard University Press, Cambridge, Mass., 1976. xiv, 490 pp., illus. $30. A Commonwealth Fund Book. Science 193, 567–568 (1976).

31. Deutsch, J. A., Moore, B. O. & Heinrichs, S. C. Unlearned specific appetite for protein. Physiology & Behavior 46, 619–624 (1989).

32. Hughes, B. O. & Wood-Gush, D. G. A specific appetite for calcium in domestic chickens. Animal Behaviour 19, 490–499 (1971).

33. Richter, C. P. & Barelare, B. NUTRITIONAL REQUIREMENTS OF PREGNANT AND LACTATING RATS STUDIED BY THE SELFSELECTION METHOD. Endocrinology 23, 15–24 (1938).

34. Leitão-Gonçalves, R. et al. Commensal bacteria and essential amino acids control food choice behavior and reproduction. PLOS Biology 15, e2000862 (2017).

35. Itskov, P. M. et al. Automated monitoring and quantitative analysis of feeding behaviour in Drosophila. Nature communications 5, 4560 (2014).

36. Carvalho-Santos, Z. et al. Cellular metabolic reprogramming controls sugar appetite in Drosophila. Nat Metab 2, 958–973 (2020).

37. Ribeiro, C. & Dickson, B. J. Sex Peptide Receptor and Neuronal TOR/S6K Signaling Modulate Nutrient Balancing in Drosophila. Current Biology 20, 1000–1005 (2010).

38. Corrales-Carvajal, V. M., Faisal, A. A. & Ribeiro, C. Internal states drive nutrient homeostasis by modulating exploration-exploitation trade-off. eLife 5, e19920 (2016).

39. Steck, K. et al. Internal amino acid state modulates yeast taste neurons to support protein homeostasis in Drosophila. eLife 7, e31625 (2018).

40. Walker, S. J., Corrales-Carvajal, V. M. & Ribeiro, C. Postmating Circuitry Modulates Salt Taste Processing to Increase Reproductive Output in Drosophila. Current Biology 25, 2621–2630 (2015).

41. Carvalho-Santos, Z. The dynamic relationship between oogenesis and whole-animal physiology. 417–449 (2026) doi:10.1016/B978-0-323-95424-2.00027-3.

42. Drummond-Barbosa, D. & Spradling, A. C. Stem cells and their progeny respond to nutritional changes during Drosophila oogenesis. Dev Biol 231, 265–78 (2001).

43. Grandison, R. C., Piper, M. D. & Partridge, L. Amino-acid imbalance explains extension of lifespan by dietary restriction in Drosophila. Nature 462, 1061–4 (2009).

44. Good, T. P. & Tatar, M. Age-specific mortality and reproduction respond to adult dietary restriction in Drosophila melanogaster. Journal of Insect Physiology 47, 1467–1473 (2001).

45. Partridge, L., Piper, M. D. W. & Mair, W. Dietary restriction in *Drosophila*. Mechanisms of Ageing and Development 126, 938–950 (2005).

46. Mirth, C. K., Nogueira Alves, A. & Piper, M. D. Turning food into eggs: insights from nutritional biology and developmental physiology of Drosophila. Current Opinion in Insect Science 31, 49–57 (2019).

47. Terashima, J. & Bownes, M. Translating Available Food Into the Number of Eggs Laid by Drosophila melanogaster. Genetics 167, 1711–1719 (2004).

48. Lushchak, O. V. et al. Specific Dietary Carbohydrates Differentially Influence the Life Span and Fecundity of Drosophila melanogaster. The Journals of Gerontology: Series A 69, 3–12 (2014).

49. Sieber, M. H., Thomsen, M. B. & Spradling, A. C. Electron Transport Chain Remodeling by GSK3 during Oogenesis Connects Nutrient State to Reproduction. Cell 164, 420–432 (2016).

50. Sieber, M. H. & Spradling, A. C. Steroid Signaling Establishes a Female Metabolic State and Regulates SREBP to Control Oocyte Lipid Accumulation. Current Biology 25, 993–1004 (2015).

51. Berg, C., Sieber, M. & Sun, J. Finishing the egg. Genetics 226, iyad183 (2024).

52. Hinnant, T. D., Merkle, J. A. & Ables, E. T. Coordinating Proliferation, Polarity, and Cell Fate in the Drosophila Female Germline. Front. Cell Dev. Biol. 8, (2020).

53. Bownes, M. & Blair, M. The effects of a sugar diet and hormones on the expression of the *Drosophila* yolk-protein genes. Journal of Insect Physiology 32, 493–501 (1986).

54. Bownes, M. & Reid, G. The role of the ovary and nutritional signals in the regulation of fat body yolk protein gene expression in *Drosophila melanogaster*. Journal of Insect Physiology 36, 471–479 (1990).

55. Pritchett, T. L., Tanner, E. A. & McCall, K. Cracking open cell death in the Drosophila ovary. Apoptosis 14, 969 (2009).

56. Shimada, Y., Burn, K. M., Niwa, R. & Cooley, L. Reversible response of protein localization and microtubule organization to nutrient stress during Drosophila early oogenesis. Dev Biol 355, 250–262 (2011).

57. Burn, K. M. et al. Somatic insulin signaling regulates a germline starvation response in Drosophila egg chambers. Developmental Biology 398, 206–217 (2015).

58. Sun, J. et al. Drosophila FIT is a protein-specific satiety hormone essential for feeding control. Nat Commun 8, 14161 (2017).

59. Zirin, J. et al. Large-Scale Transgenic Drosophila Resource Collections for Loss- and Gain-of-Function Studies. Genetics 214, 755–767 (2020).

60. Hu, Y. et al. FlyRNAi.org—the database of the *Drosophila* RNAi screening center and transgenic RNAi project: 2017 update. Nucleic Acids Res 45, D672–D678 (2017).

61. Dietzl, G. et al. A genome-wide transgenic RNAi library for conditional gene inactivation in Drosophila. Nature 448, 151–156 (2007).

62. Heredia, F. et al. The steroid-hormone ecdysone coordinates parallel pupariation neuromotor and morphogenetic subprograms via epidermis-to-neuron Dilp8-Lgr3 signal induction. Nat Commun 12, 3328 (2021).

63. Mackay, T. F. C. et al. The Drosophila melanogaster Genetic Reference Panel. Nature 482, 173–178 (2012).

64. Öztürk-Çolak, A. et al. FlyBase: updates to the Drosophila genes and genomes database. Genetics 227, iyad211 (2024).

65. Jenkins, V. K., Larkin, A. & Thurmond, J. Using FlyBase: A Database of Drosophila Genes and Genetics. in Drosophila: Methods and Protocols (ed. Dahmann, C.) 1–34 (Springer US, New York, NY, 2022). doi:10.1007/978-1-0716-2541-5_1.

66. Deshpande, S. A. et al. Quantifying Drosophila food intake: comparative analysis of current methodology. Nat Methods 11, 535–540 (2014).

67. Schindelin, J., et al. Fiji: an open-source platform for biological-image analysis. Nat Methods 9, 676–682 (2012).

68. Jia, D., Xu, Q., Xie, Q., Mio, W. & Deng, W.-M. Automatic stage identification of Drosophila egg chamber based on DAPI images. Sci Rep 6, 18850 (2016).

69. Yan, D. et al. A Regulatory Network of Drosophila Germline Stem Cell Self-Renewal. Developmental Cell 28, 459–473 (2014).

70. Carvalho, G. B., Kapahi, P., Anderson, D. J. & Benzer, S. Allocrine Modulation of Feeding Behavior by the Sex Peptide of Drosophila. Current Biology 16, 692–696 (2006).

71. Toshima, N. & Tanimura, T. Taste preference for amino acids is dependent on internal nutritional state in Drosophila melanogaster. J Exp Biol 215, 2827–2832 (2012).

72. Soller, M., Bownes, M. & Kubli, E. Control of Oocyte Maturation in Sexually MatureDrosophilaFemales. Developmental Biology 208, 337–351 (1999).

73. Heifetz, Y., Lung, O., Frongillo, E. A. & Wolfner, M. F. The *Drosophila* seminal fluid protein Acp26Aa stimulates release of oocytes by the ovary. Current Biology 10, 99–102 (2000).

74. Herndon, L. A. & Wolfner, M. F. A Drosophila seminal fluid protein, Acp26Aa, stimulates egg laying in females for 1 day after mating. Proceedings of the National Academy of Sciences 92, 10114–10118 (1995).

75. Liu, H. & Kubli, E. Sex-peptide is the molecular basis of the sperm effect in Drosophila melanogaster. Proceedings of the National Academy of Sciences 100, 9929–9933 (2003).

76. Yapici, N., Kim, Y.-J., Ribeiro, C. & Dickson, B. J. A receptor that mediates the post-mating switch in Drosophila reproductive behaviour. Nature 451, 33–37 (2008).

77. Wang, F. et al. Neural circuitry linking mating and egg laying in Drosophila females. Nature 579, 101–105 (2020).

78. Wolfner, M. F. The gifts that keep on giving: physiological functions and evolutionary dynamics of male seminal proteins in Drosophila. Heredity 88, 85–93 (2002).

79. Gillott, C. Male Accessory Gland Secretions: Modulators of Female Reproductive Physiology and Behavior. Annu. Rev. Entomol. 48, 163–184 (2003).

80. Chapman, T. & Davies, S. J. Functions and analysis of the seminal fluid proteins of male Drosophila melanogaster fruit flies. Peptides 25, 1477–1490 (2004).

81. Ravi Ram, K. & Wolfner, M. F. Seminal influences: Drosophila Acps and the molecular interplay between males and females during reproduction. Integrative and Comparative Biology 47, 427–445 (2007).

82. Aigaki, T., Fleischmann, I., Chen, P.-S. & Kubli, E. Ectopic expression of sex peptide alters reproductive behavior of female D. melanogaster. Neuron 7, 557–563 (1991).

83. Häsemeyer, M., Yapici, N., Heberlein, U. & Dickson, B. J. Sensory Neurons in the *Drosophila* Genital Tract Regulate Female Reproductive Behavior. Neuron 61, 511–518 (2009).

84. Yang, C. et al. Control of the Postmating Behavioral Switch in Drosophila Females by Internal Sensory Neurons. Neuron 61, 519–526 (2009).

85. Rezával, C. et al. Neural Circuitry Underlying Drosophila Female Postmating Behavioral Responses. Current Biology 22, 1155–1165 (2012).

86. Feng, K., Palfreyman, M. T., Häsemeyer, M., Talsma, A. & Dickson, B. J. Ascending SAG Neurons Control Sexual Receptivity of *Drosophila* Females. Neuron 83, 135–148 (2014).

87. Guss, J. L. & Kissileff, H. R. Microstructural analyses of human ingestive patterns: from description to mechanistic hypotheses. Neuroscience & Biobehavioral Reviews 24, 261–268 (2000).

88. Davis, J. D. & Smith, G. P. Analysis of the microstructure of the rhythmic tongue movements of rats ingesting maltose and sucrose solutions. Behavioral Neuroscience 106, 217–228 (1992).

89. Davis, J. D. & Levine, M. W. A model for the control of ingestion. Psychological Review 84, 379–412 (1977).

90. Davis, J. D. & Perez, M. C. Food deprivation- and palatability-induced microstructural changes in ingestive behavior. *American Journal of Physiology-Regulatory*, Integrative and Comparative Physiology 264, R97–R103 (1993).

91. Münch, D., Goldschmidt, D. & Ribeiro, C. The neuronal logic of how internal states control food choice. Nature 607, 747–755 (2022).

92. Tastekin, I. et al. Connectomics Reveals a Feed-Forward Swallowing Circuit Driving Protein Appetite. 2025.08.25.671815 Preprint at 10.1101/2025.08.25.671815 (2025).

93. Berg, C., Sieber, M. & Sun, J. Finishing the egg. Genetics 226, iyad183 (2024).

94. Berleth, T. et al. The role of localization of bicoid RNA in organizing the anterior pattern of the Drosophila embryo. EMBO J 7, 1749–1756 (1988).

95. Cox, D. N., Seyfried, S. A., Jan, L. Y. & Jan, Y. N. Bazooka and atypical protein kinase C are required to regulate oocyte differentiation in the Drosophila ovary. Proceedings of the National Academy of Sciences 98, 14475–14480 (2001).

96. Johnston, D. St., Driever, W., Berleth, T., Richstein, S. & Nüsslein-Volhard, C. Multiple steps in the localization of bicoid RNA to the anterior pole of the Drosophila oocyte. Development 107, 13–19 (1989).

97. Kim, S. et al. Kinase-activity-independent functions of atypical protein kinase C in Drosophila. J Cell Sci 122, 3759–3771 (2009).

98. Lasko, P. Patterning the Drosophila embryo: A paradigm for RNA-based developmental genetic regulation. WIREs RNA 11, e1610 (2020).

99. Llamazares, S., Tavosanis, G. & Gonzalez, C. Cytological characterisation of the mutant phenotypes produced during early embryogenesis by null and loss-of-function alleles of the γTub37C gene in Drosophila. J Cell Sci 112, 659–667 (1999).

100. Morisato, D. & Anderson, K. V. The *spätzle* gene encodes a component of the extracellular signaling pathway establishing the dorsal-ventral pattern of the Drosophila embryo. Cell 76, 677–688 (1994).

101. Schneider, D. S., Jin, Y., Morisato, D. & Anderson, K. V. A processed form of the Spätzle protein defines dorsal-ventral polarity in the Drosophila embryo. Development 120, 1243–1250 (1994).

102. Schnorrer, F., Luschnig, S., Koch, I. & Nüsslein-Volhard, C. *γ-Tubulin37C* and *γ-tubulin ring complex protein 75* Are Essential for *bicoid* RNA Localization during *Drosophila* Oogenesis. Developmental Cell 3, 685–696 (2002).

103. Stein, D. S. & Stevens, L. M. Maternal control of the Drosophila dorsal–ventral body axis. WIREs Developmental Biology 3, 301–330 (2014).

104. Tavosanis, G., Llamazares, S., Goulielmos, G. & Gonzalez, C. Essential role for γ-tubulin in the acentriolar female meiotic spindle of Drosophila. EMBO J 16, 1809–1819 (1997).

105. Tian, A.-G. & Deng, W.-M. Lgl and its phosphorylation by aPKC regulate oocyte polarity formation in Drosophila. Development 135, 463–471 (2008).

106. Weil, T. T., Parton, R., Davis, I. & Gavis, E. R. Changes in *bicoid* mRNA Anchoring Highlight Conserved Mechanisms during the Oocyte-to-Embryo Transition. Current Biology 18, 1055–1061 (2008).

107. Garelli, A., Gontijo, A. M., Miguela, V., Caparros, E. & Dominguez, M. Imaginal Discs Secrete Insulin-Like Peptide 8 to Mediate Plasticity of Growth and Maturation. Science 336, 579–582 (2012).

108. Colombani, J., Andersen, D. S. & Léopold, P. Secreted Peptide Dilp8 Coordinates Drosophila Tissue Growth with Developmental Timing. Science 336, 582–585 (2012).

109. Garelli, A. et al. Dilp8 requires the neuronal relaxin receptor Lgr3 to couple growth to developmental timing. Nat Commun 6, 8732 (2015).

110. Vallejo, D. M., Juarez-Carreño, S., Bolivar, J., Morante, J. & Dominguez, M. A brain circuit that synchronizes growth and maturation revealed through Dilp8 binding to Lgr3. Science 350, aac6767 (2015).

111. Boulan, L., Andersen, D., Colombani, J., Boone, E. & Léopold, P. Inter-Organ Growth Coordination Is Mediated by the Xrp1-Dilp8 Axis in Drosophila. Developmental Cell 49, 811–818.e4 (2019).

112. Gontijo, A. M. & Garelli, A. The biology and evolution of the Dilp8-Lgr3 pathway: A relaxin-like pathway coupling tissue growth and developmental timing control. Mechanisms of Development 154, 44–50 (2018).

113. Blanco-Obregon, D. et al. A Dilp8-dependent time window ensures tissue size adjustment in Drosophila. Nat Commun 13, 5629 (2022).

114. Oramas, R., Yacuk, K., Cho, S. E., Aloisio, N. R. & Sun, J. ILP8 serves as a mature follicle sensor to prevent excessive accumulation of mature follicles in Drosophila ovaries and oocyte aging. Development 152, dev204827 (2025).

115. Volonté, Y. et al. Dilp8 relaxin signaling from ovarian follicle cells to Lgr3+ neurons promotes spontaneous ovulation and oocyte quality in Drosophila. Development dev.204955 (2026) doi:10.1242/dev.204955.

116. Heigwer, F., Port, F. & Boutros, M. RNA Interference (RNAi) Screening in Drosophila. Genetics 208, 853–874 (2018).

117. Mehta, S. J. K., Joshi, P. A. & Mishra, R. K. Molecular and Phenotypic Characterization Following RNAi Mediated Knockdown in *Drosophila*. Bio-protocol 11, (2021).

118. López-Varea, A. et al. Genome-wide phenotypic RNAi screen in the *Drosophila* wing: global parameters. G3 Genes|Genomes|Genetics 11, jkab351 (2021).

119. Laturney, M., Sterne, G. R. & Scott, K. Mating activates neuroendocrine pathways signaling hunger in Drosophila females. eLife 12, e85117 (2023).

120. Carvalho-Santos, Z. & Ribeiro, C. Gonadal ecdysone titers are modulated by protein availability but do not impact protein appetite. J Insect Physiol 106, 30–35 (2018).

121. Ables, E. T. & Drummond-Barbosa, D. Steroid Hormones and the Physiological Regulation of Tissue-Resident Stem Cells: Lessons from the Drosophila Ovary. Curr Stem Cell Rep 3, 9–18 (2017).

122. Morris, L. X. & Spradling, A. C. Steroid Signaling within Drosophila Ovarian Epithelial Cells Sex-Specifically Modulates Early Germ Cell Development and Meiotic Entry. PLoS ONE 7, e46109 (2012).

123. Bownes, M., Ronaldson, E. & Mauchline, D. 20-Hydroxyecdysone, but Not Juvenile Hormone, Regulation of*yolk protein*Gene Expression Can Be Mapped to*cis*-Acting DNA Sequences. Developmental Biology 173, 475–489 (1996).

124. Ables, E. T., Hwang, G. H., Finger, D. S., Hinnant, T. D. & Drummond-Barbosa, D. A Genetic Mosaic Screen Reveals Ecdysone-Responsive Genes Regulating Drosophila Oogenesis. G3 Genes|Genomes|Genetics 6, 2629–2642 (2016).

125. Knapp, E. & Sun, J. Steroid signaling in mature follicles is important for Drosophila ovulation. Proceedings of the National Academy of Sciences 114, 699–704 (2017).

126. Akyaw, P. A., Paulo, T. F., Lafuente, E. & Sucena, É. Pathogen-induced damage in Drosophila: Uncoupling disease tolerance from resistance. PLOS Pathogens 21, e1013482 (2025).

127. Babin, A., Gatti, J.-L. & Poirié, M. Bacillus thuringiensis bioinsecticide influences Drosophila oviposition decision. R Soc Open Sci. 10, 230565 (2023).

128. Billeter, J.-C. & Wolfner, M. F. Chemical Cues that Guide Female Reproduction in Drosophila melanogaster. J Chem Ecol 44, 750–769 (2018).

129. Bixler, A. & Schnee, F. B. The effects of the timing of exposure to cadmium on the oviposition behavior of Drosophila melanogaster. Biometals 31, 1075–1080 (2018).

130. Finetti, L. et al. Monarda didyma Hydrolate Affects the Survival and the Behaviour of Drosophila suzukii. Insects 13, (2022).

131. Fowler, E. K. et al. Female oviposition decisions are influenced by the microbial environment. *j. evol*. Biol. 38, 379–390 (2025).

132. Joseph, R. M., Devineni, A. V., King, I. F. G. & Heberlein, U. Oviposition preference for and positional avoidance of acetic acid provide a model for competing behavioral drives in Drosophila. Proceedings of the National Academy of Sciences 106, 11352–11357 (2009).

133. Li, Y. et al. Capsaicin Functions as Drosophila Ovipositional Repellent and Causes Intestinal Dysplasia. Sci Rep 10, 9963 (2020).

134. Liu, Z.-H. et al. Oxidative stress caused by lead (Pb) induces iron deficiency in *Drosophila melanogaster*. Chemosphere 243, 125428 (2020).

135. Liu, Z.-H. et al. Mitochondrial iron deficiency mediated inhibition of ecdysone synthesis underlies lead (Pb) induced developmental toxicity in *Drosophila melanogaster*. Toxicology and Applied Pharmacology 497, 117283 (2025).

136. Mansourian, S. et al. Fecal-Derived Phenol Induces Egg-Laying Aversion in *Drosophila*. Current Biology 26, 2762–2769 (2016).

137. Asarian, L. & Geary, N. Modulation of appetite by gonadal steroid hormones. Philos Trans R Soc Lond B Biol Sci 361, 1251–1263 (2006).

138. Butera, P. C. Estradiol and the control of food intake. Physiology & Behavior 99, 175–180 (2010).

139. Candan, E., Metin, Z. E. & Tengilimoglu-Metin, M. M. The role of premenstrual syndrome in hedonic hunger and food craving during the menstrual cycle. J Nutr Sci 14, e66 (2025).

140. Dye, L. & Blundell, J. E. Menstrual cycle and appetite control: implications for weight regulation. Human Reproduction 12, 1142–1151 (1997).

141. González-García, I. & Xu, Y. Hypothalamic actions of estrogens in the regulation of energy and glucose homeostasis. Rev Endocr Metab Disord 10.1007/s11154-025-09994-1 (2025) doi:10.1007/s11154-025-09994-1.

142. Hirschberg, A. L. Sex hormones, appetite and eating behaviour in women. Maturitas 71, 248–256 (2012).

143. Nijboer, A. C. S., Sellitto, M., Ruitenberg, M. F. L., Kerkkänen, K. I. L. & Schomaker, J. Food-related exploration across the menstrual cycle. Appetite 196, 107261 (2024).

144. Santollo, J. & Daniels, D. Multiple estrogen receptor subtypes influence ingestive behavior in female rodents. Physiology & Behavior 152, 431–437 (2015).

145. Zhu, J., Zhou, Y., Jin, B. & Shu, J. Role of estrogen in the regulation of central and peripheral energy homeostasis: from a menopausal perspective. Therapeutic Advances in Endocrinology 14, 20420188231199359 (2023).

