## Supplemental Figures and Tables for "Ovary-Derived Signals Align Protein Appetite with Oogenesis"

[3] iNOVA4Health, Nova Medical School, Faculdade de Ciências Médicas, NMS, FCM, Universidade Nova de Lisboa, Lisbon, Portugal

[4] Behavior and Metabolism Laboratory, Champalimaud Research, Champalimaud Foundation, Lisbon, Portugal

[5] FMUL – Faculdade de Medicina da Universidade de Lisboa, Lisbon, Portugal

### Supplementary Figures

### Nóbrega, R. R. *et al.* - Supplementary Figure 1

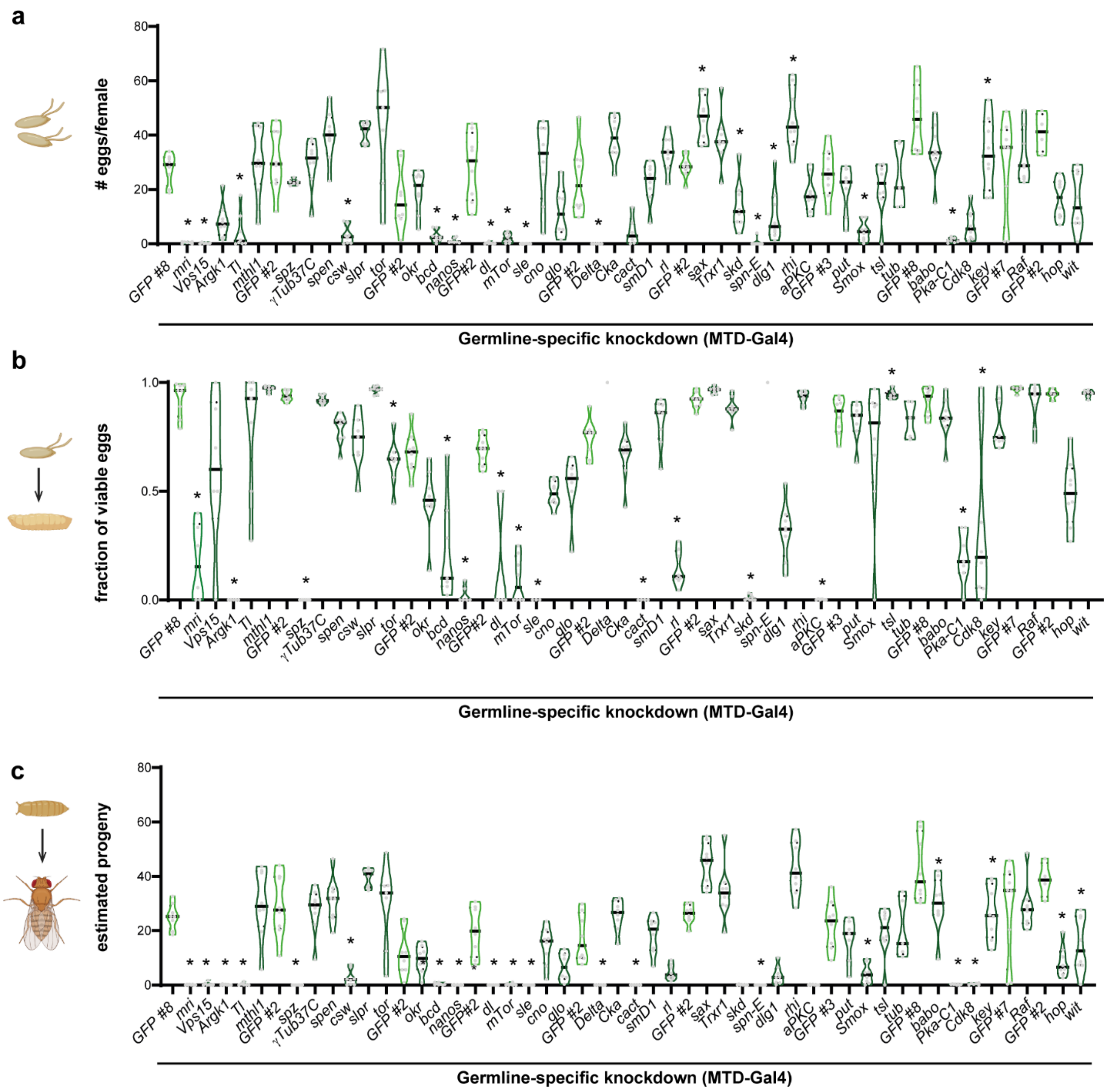


**Supplementary Figure 1 – Germline-specific gene knockdowns lead to decreased female fertility.** Fully fed mated females knocked down for the indicated genes using the MTD-Gal4 driver were assayed for fertility. **a** Average number of eggs laid per female in 24 hours; **b** average fraction of viable eggs per female and **c** estimated progeny. **a-c** Data are represented as truncated violin plots and median and quartiles are represented in solid or dashed lines, respectively. *GFP* RNAi was used as negative control (GFP#8 – valium 22, attP2; GFP#2 - valium 20; attP2; GFP#3 - valium 20; attp40; GFP#7 – valium 22, attP40). n = 4-8. The statistical significance was determined using ordinary one-way ANOVA or t-test in cases where the data followed a normal distribution, and the Kruskal-Wallis test or Mann-Whitney in the remaining cases. Significance is marked with * $p<0.05$. The detailed p-values are shown in **Supplementary Table 6**.

**Nóbrega, R. R. *et al.* - Supplementary Figure 2**

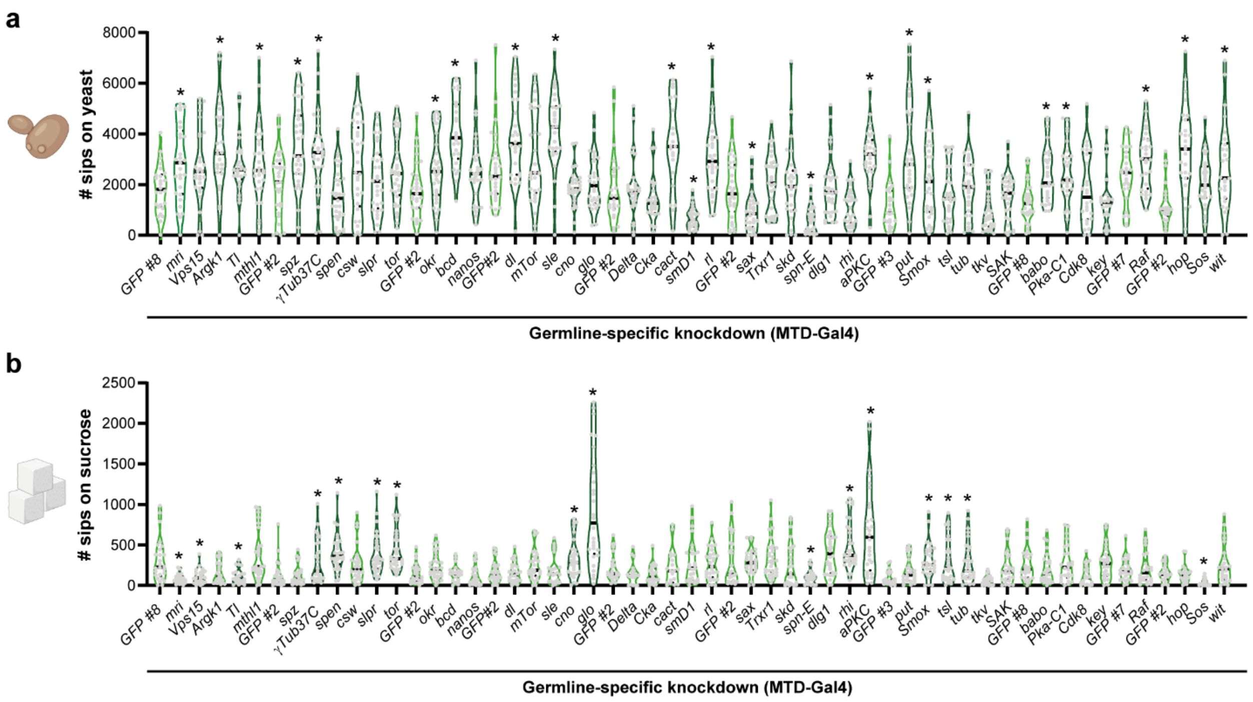


**Supplementary Figure 2 – Ovarian physiology impacts nutrient appetite.** Fully fed mated females knocked down for the indicated genes using the MTD-Gal4 driver were assayed for feeding behavior using the flyPAD. Total number of sips on yeast **(a)** or sucrose **(b)** are represented as truncated violin plots and each dot corresponds to one individual fly. Median and quartiles are represented in solid or dashed lines, respectively. *GFP* RNAi was used as negative control (GFP#8 – valium 22, attP2; GFP#2 - valium 20; attP2; GFP#3 - valium 20; attp40; GFP#7 – valium 22, attP40). n = 19-32. The statistical significance was determined using ordinary one-way ANOVA or t-test in cases where the data followed a normal distribution, and the Kruskal-Wallis test or Mann-Whitney in the remaining cases. Significance is marked with * p<0.05. The detailed p-values are shown in **Supplementary Table 7**.

### Nóbrega, R. R. *et al.* - Supplementary Figure 3

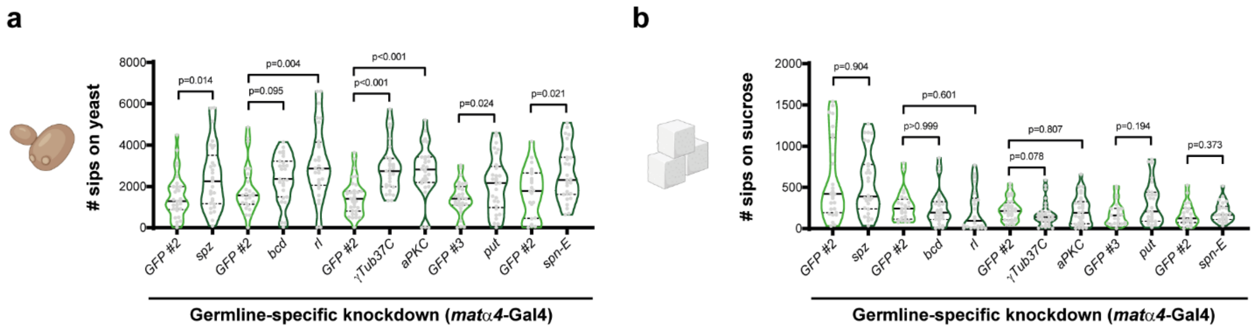


**Supplementary Figure 3 - Feeding effects in germline-manipulated females arise from disruptions to later oogenesis. a, b** Fully fed mated females knocked down for the indicated genes in the germline using the *matα*-Gal4 driver were assayed for feeding behavior using the flyPAD. Total number of sips on yeast **(a)** or sucrose **(b)** are represented as truncated violin plots and each dot corresponds to one individual fly. Median and quartiles are represented in solid or dashed lines, respectively. *GFP* RNAi was used as negative control (GFP#2 - valium 20; attP2; GFP#3 - valium 20; attp40). n = 22-32. The statistical significance was determined using ordinary one-way ANOVA or t-test in cases where the data followed a normal distribution, and the Kruskal-Wallis test or Mann-Whitney in the remaining cases.

### Nóbrega, R. R. *et al.* - Supplementary Figure 4

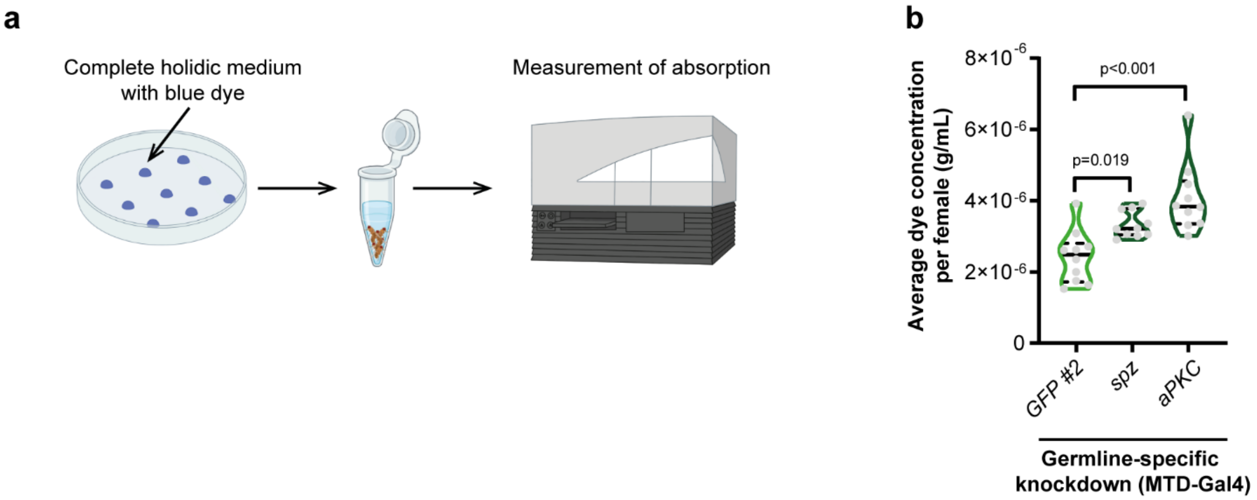


**Supplementary Figure 4 – Ovarian physiology increases appetite for nutrient-rich foods. a** Schematic representation of colorimetric feeding assay. **b** Fully fed mated females knocked down for the indicated genes using the MTD-Gal4 driver were assayed for feeding behavior using the Colorimetric feeding assay. *GFP* RNAi was used as negative control (GFP#2 - valium 20; attP2). n = 10. Data points are represented as truncated violin plots and median and quartiles are represented in solid or dashed lines, respectively. The statistical significance was determined using ordinary one-way ANOVA.

Nóbrega, R. R. *et al.* - Supplementary Figure 5

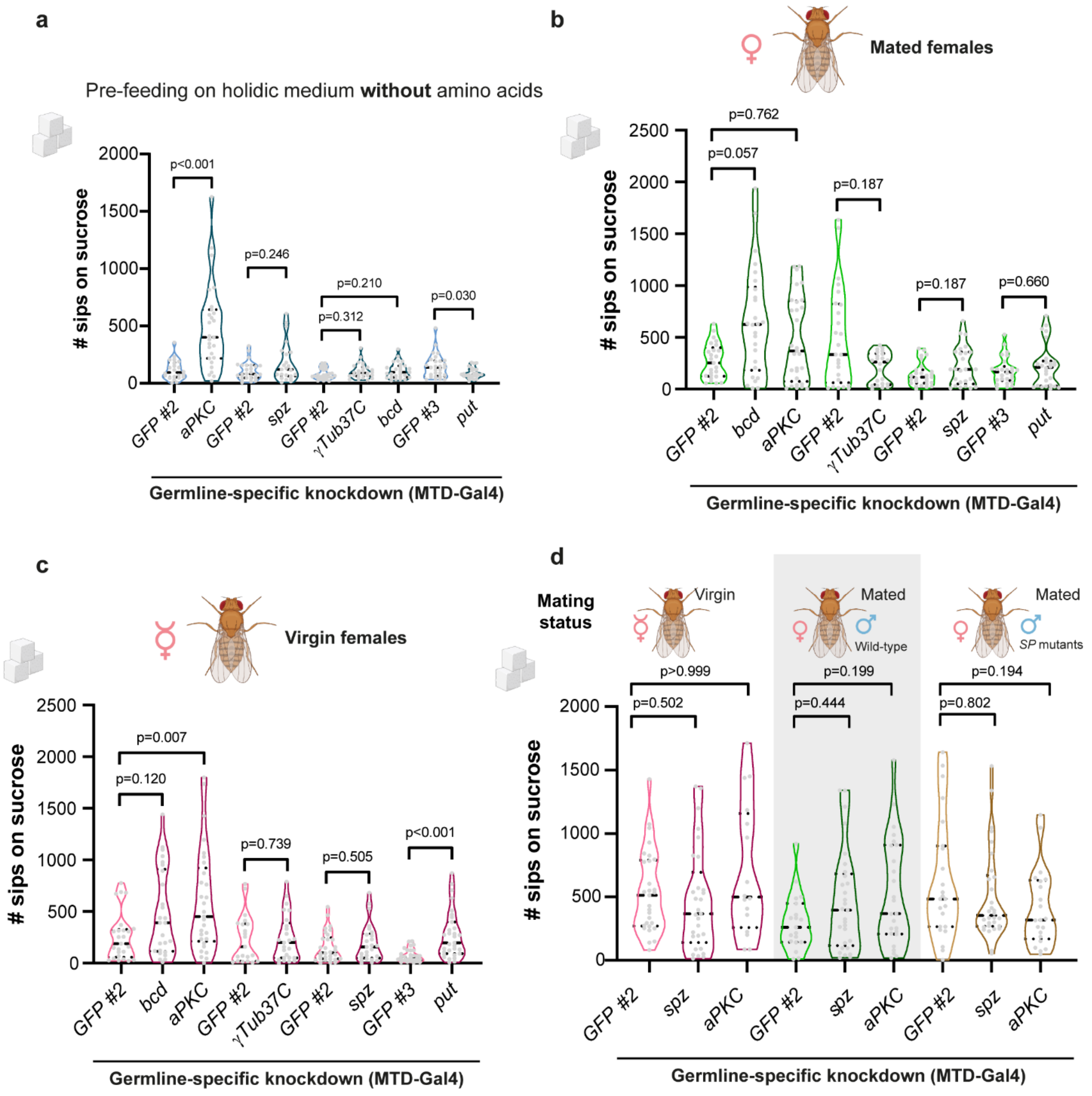


**Supplementary Figure 5 – Sucrose appetite is not consistently affected by ovarian physiology. a** Females knocked down for the indicated genes in the germline using the MTD-Gal4 driver were pre-treated for 4 days with a synthetic diet from where all amino acids were omitted. The females were assayed for feeding behavior using the flyPAD. n = 25-28. **b-d** Fully fed females mated with wild type (**b**) or Sex Peptide (SP) mutant males **(d)** or virgins **(c, d)** knocked down for the indicated genes in the germline using the MTD-Gal4 driver were assayed for feeding behavior using the flyPAD. **a-d** Total number of sips on sucrose are represented as truncated violin plots and each dot corresponds to one individual fly. Median and quartiles are represented in solid or dashed lines, respectively. *GFP* RNAi was used as negative control (GFP#2 - valium 20; attP2; GFP#3 - valium 20; attp40). n = 21-34. The statistical significance was determined using ordinary one-way ANOVA or t-test in cases where the data followed a normal distribution, and the Kruskal-Wallis test or Mann-Whitney in the remaining cases.

**Nóbrega, R. R. *et al.* - Supplementary Figure 6**

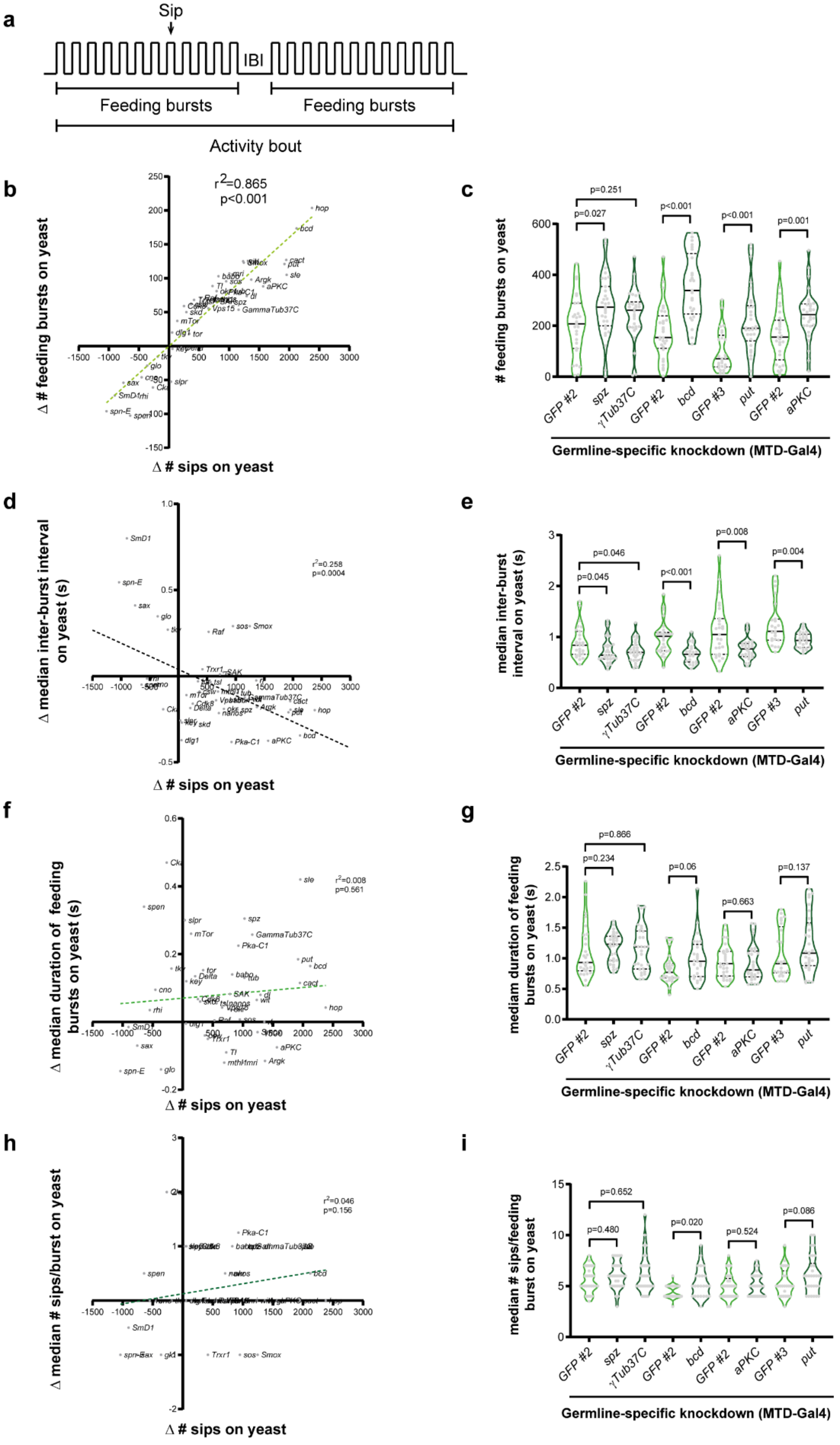

**Supplementary Figure 6 – Ovarian physiology regulates yeast appetite by modulating the number of feeding bursts. a** Illustration of the feeding temporal dynamics of a fly on a gelatinous food, depicting the different parameters measured and analyzed using the flyPAD behavioral setup[35]. IBI – Inter-burst interval. **b, d, f, h** Correlation between the number of sips and the number of feeding bursts (**b**), the median duration of feeding bursts (**d**), the median number of sips per burst (**f**) or the median inter-burst interval (**h**) for each genetic manipulation. For further details, see Materials and Methods. The regression line represents the best fit for the data. **c, e, g, I** Fully fed mated females knocked down for the indicated genes in the germline were assayed for feeding behavior using the flyPAD. The number of feeding bursts (**c**), the median duration of feeding bursts (**e**), the median number of sips per burst (**g**) and the median inter-burst interval (**I**) are represented as truncated violin plots and each dot corresponds to one individual fly. Median and quartiles are represented in solid or dashed lines, respectively. *GFP* RNAi was used as negative control (GFP#2 - valium 20; attP2; GFP#3 - valium 20; attp40). n = 25-31. The statistical significance was determined using ordinary one-way ANOVA or t-test in cases where the data followed a normal distribution, and the Kruskal-Wallis test or Mann-Whitney in the remaining cases.

**Nóbrega, R. R. *et al.* - Supplementary Figure 7**

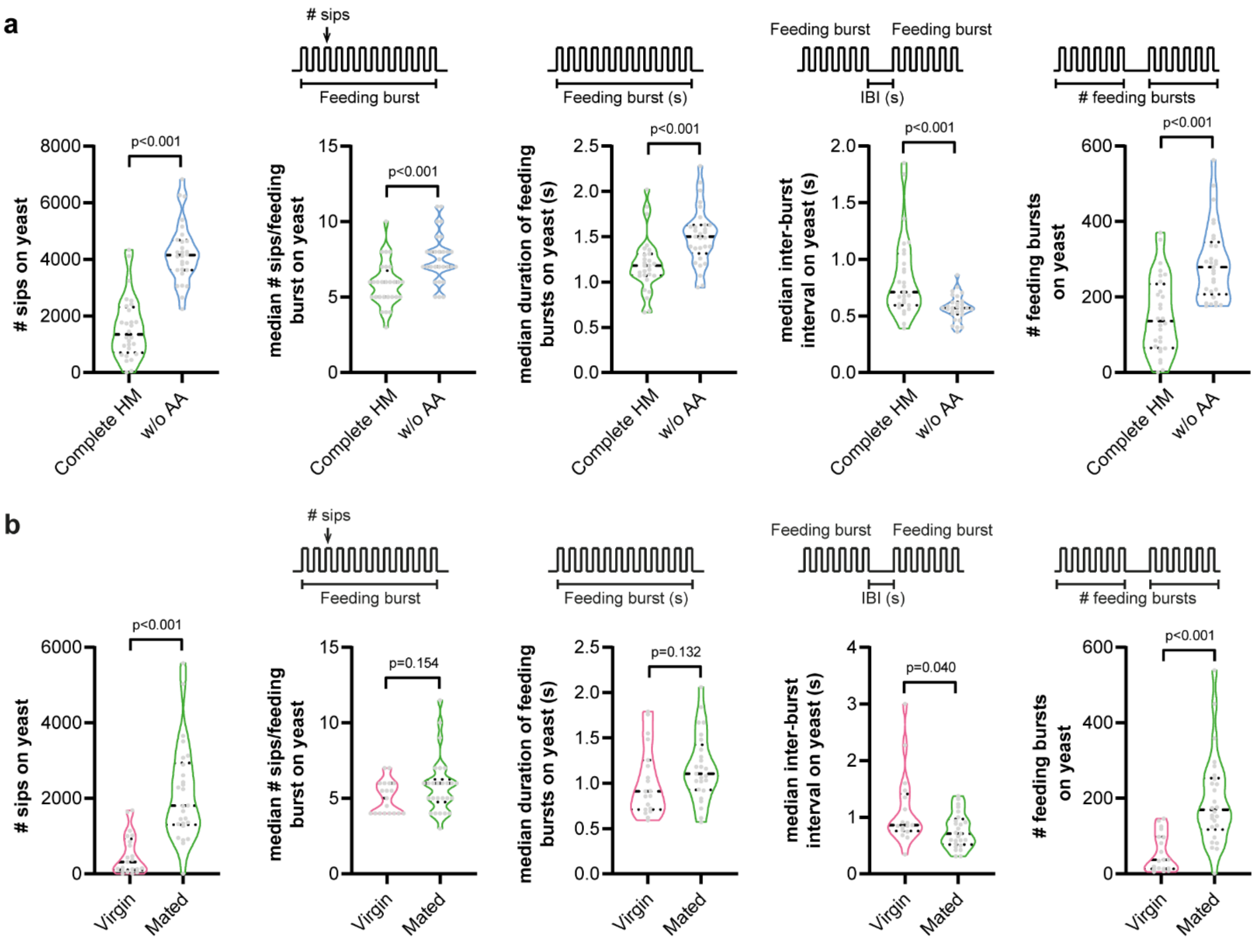


**Supplementary Figure 7 – Amino acid deprivation and mating state modulate yeast appetite through distinct feeding parameters. a** Wild-type mated females were pre-treated for 4 days with either a complete holidic medium, or a holidic medium from where all amino acids were omitted (w/o AA). Females were assayed for feeding behavior using the flyPAD. n = 27-29. Total number of sips, median number of sips per feeding burst, median feeding burst duration, median inter-burst interval and number of feeding bursts on yeast are shown from left to right. n = 22-31. **b** Wild-type fully fed virgin or mated females were assayed for feeding behavior using the flyPAD. Total number of sips, median number of sips per feeding burst, median feeding burst duration, median inter-burst interval and number of feeding bursts on yeast are shown from left to right. n=17-26. **a, b** Data are represented as truncated violin plots and each dot corresponds to one individual fly. Median and quartiles are represented in solid or dashed lines, respectively. The statistical significance was determined using t-test in cases where the data followed a normal distribution or Mann-Whitney in the remaining cases.

### Nóbrega, R. R. *et al.* - Supplementary Figure 8

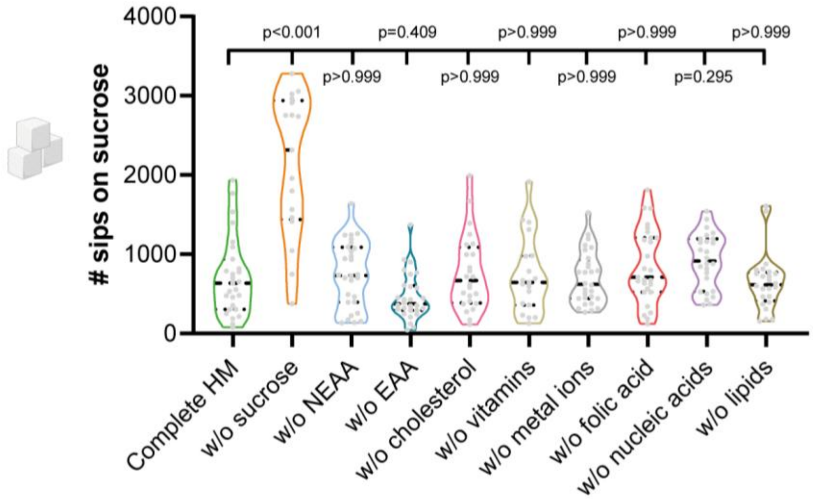


**Supplementary Figure 8 – Sucrose appetite of females fed on diets lacking specific nutrient groups. a** Wild-type mated females were pre-treated for 4 days with either a complete holidic medium, or a holidic medium from where specific groups of nutrients were omitted (w/o – without). Complete holidic medium is used as negative control. Females were assayed for feeding behavior using the flyPAD. n = 17-28. Total number of sips on sucrose are represented as truncated violin plots and each dot corresponds to one individual fly. Median and quartiles are represented in solid or dashed lines, respectively. The statistical significance was determined using the Kruskal-Wallis test.

Nóbrega, R. R. *et al.* - Supplementary Figure 9

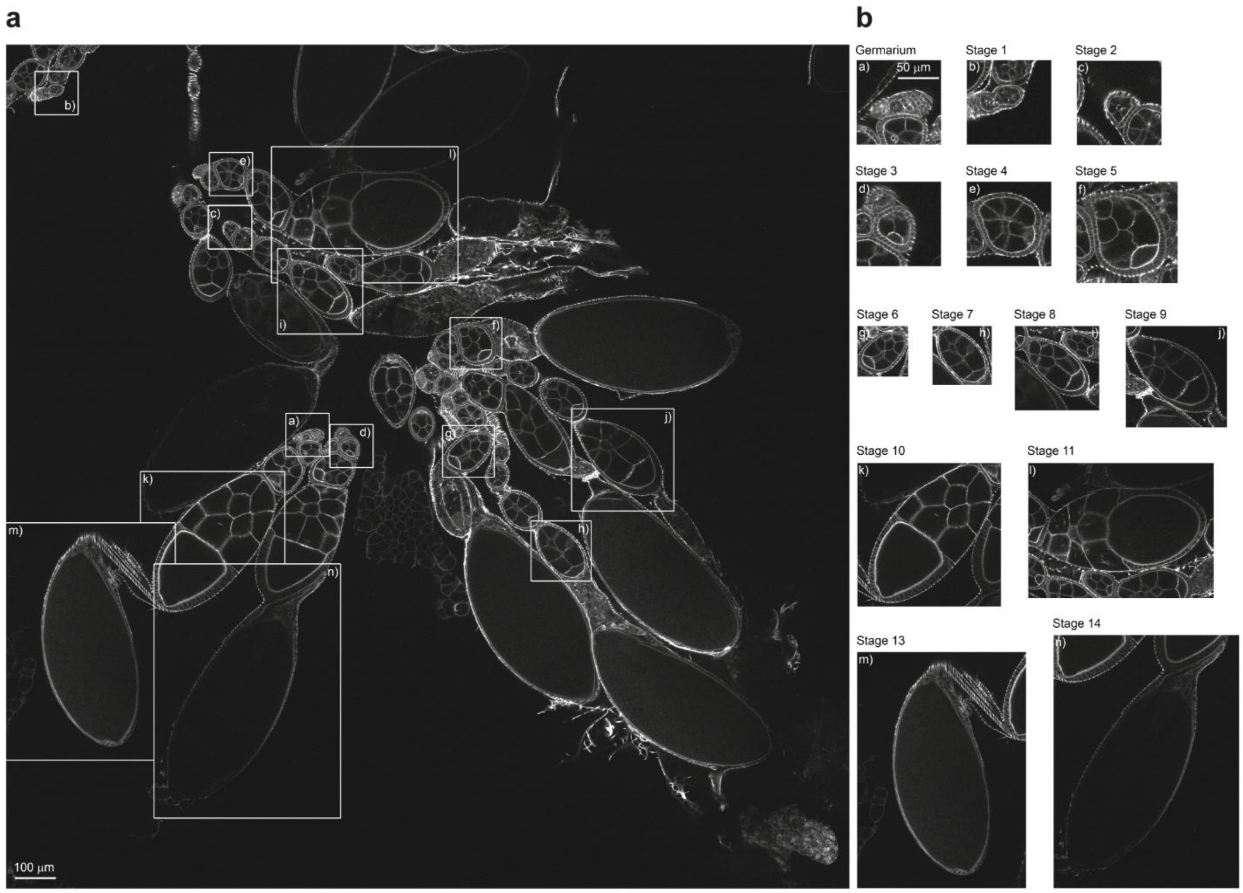


**Supplementary Figure 9 – Quantification of oogenesis stages. a** Representative image illustrating the criteria used for egg chamber stage quantification. Ovariole morphology is visualized by phalloidin staining of the actin cytoskeleton. Representative stages are indicated with insets and shown individually in **b**. Staging criteria were based on [68]. Scale bar, 100 µm. **b** Germarium to stage 5 egg chambers are shown at 2× magnification (scale bar, 50 µm). Stages 6–14 are shown at their original magnification.

### Nóbrega, R. R. *et al.* - Supplementary Figure 10

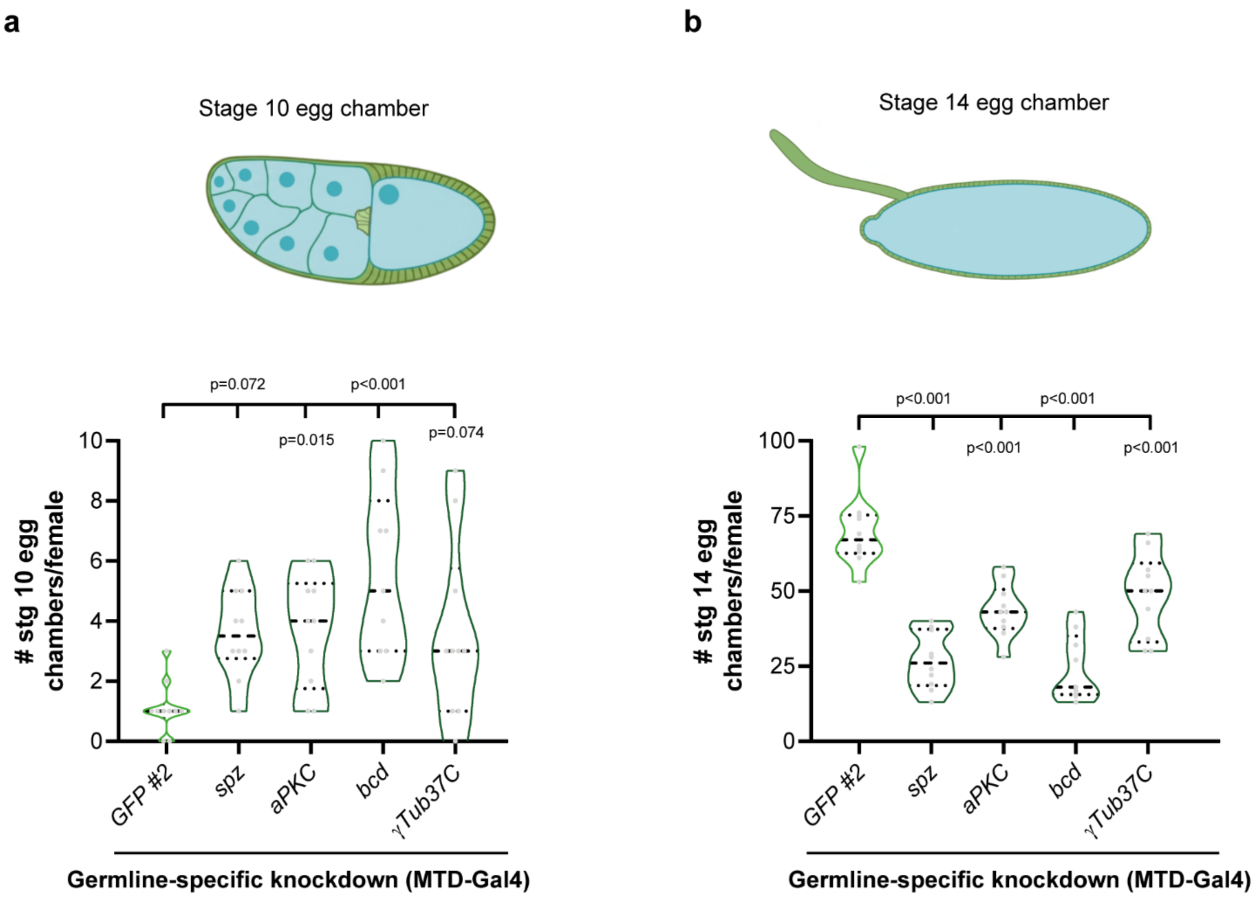


**Supplementary Figure 10 – Germline perturbations that affect yeast appetite commonly alter oogenesis progression.** Fully fed mated females knocked down for the indicated genes using the MTD-Gal4 driver were assayed for the number of stage 10 (**a**) or stage 14 (**b**) egg chambers, as determined by Phalloidin staining which reveals the actin cytoskeleton. Total number of egg chambers per stage per female are represented as truncated violin plots and median and quartiles are represented in solid or dashed lines, respectively. *GFP* RNAi was used as negative control (GFP#2 - valium 20; attP2). n = 9-10. The statistical significance was determined using ordinary one-way ANOVA in cases where the data followed a normal distribution and the Kruskal-Wallis test in the remaining cases.

**Nóbrega, R. R. *et al.* - Supplementary Figure 11**

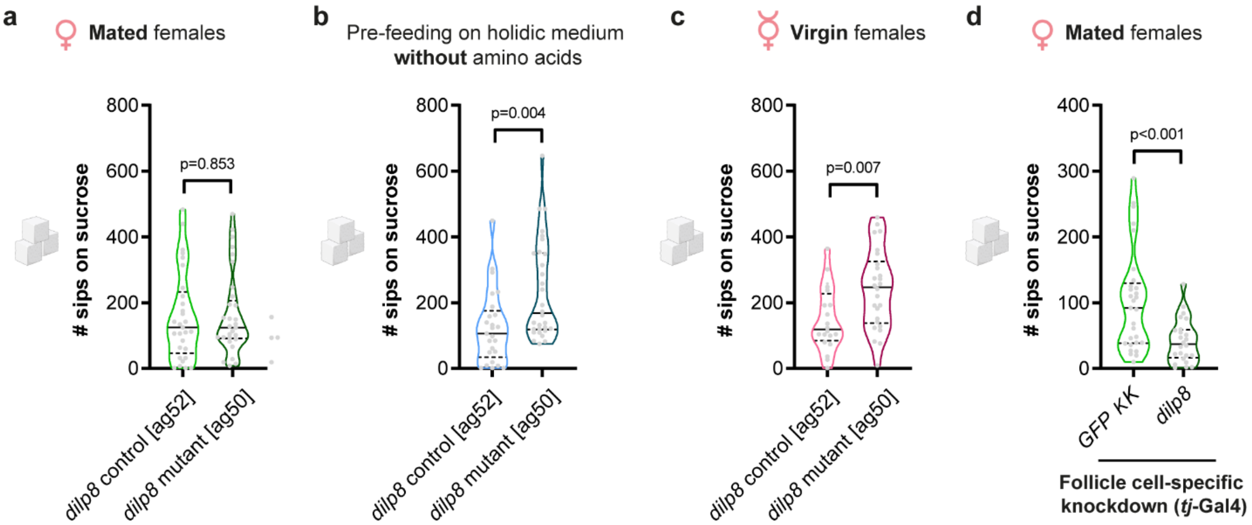


**Supplementary Figure 11 – Sucrose appetite is not consistently affected by Dilp8.** *dilp8* control or knockout mated fully fed **(a)**, mated amino-acid deprived **(b)**, or fully fed virgins **(c)**; or females knocked down for *dilp8* in the follicle cells **(d)** were assayed for feeding behavior using the flyPAD. **a-d** Total number of sips on sucrose are represented as truncated violin plots and each dot corresponds to one individual fly. Median and quartiles are represented in solid or dashed lines, respectively. The corresponding genetic background control was used as control for *Dilp8* knockout mutant **(a-c)** and *GFP* RNAi was used as negative control **(d)**. n = 20-33. The statistical significance was determined using t-test in cases where the data followed a normal distribution, and Mann-Whitney in the remaining cases.

### Supplementary Table 1 – List of stocks used in the manuscript.

| Stock name | Gene affected | Origin | Stock number | Identifier for publication |
|---|---|---|---|---|
| WT | | BDSC | 28196 | RRID:BDSC_28196 |
| MTD-Gal4 | | BDSC | 31777 | RRID:BDSC_31777 |
| Matαtub-Gal4 | | BDSC | 7063 | RRID:BDSC_7063 |
| tj-Gal4 | | Gontijo Lab | NA | NA |
| dilp8 control [ag52] | Drosophila Insulin-like peptide 8 | Gontijo Lab | NA | NA |
| dilp8 mutant [ag50] | Drosophila Insulin-like peptide 8 | Gontijo Lab | NA | NA |
| GFP #2 | | BDSC | 41553 | RRID:BDSC_41553 |
| GFP #3 | | BDSC | 41552 | RRID:BDSC_41552 |
| GFP #7 | | BDSC | 41557 | RRID:BDSC_41557 |
| GFP #8 | | BDSC | 41558 | RRID:BDSC_41558 |
| GFP KK | | VDRC | 60103 | NA |
| Argk1 | Arginine kinase 1 | BDSC | 35221 | RRID:BDSC_35221 |
| aPKC | atypical protein kinase C | BDSC | 34332 | RRID:BDSC_34332 |
| babo | baboon | BDSC | 41585 | RRID:BDSC_41585 |
| bcd | bicoid | BDSC | 33886 | RRID:BDSC_33886 |
| cact | cactus | BDSC | 34775 | RRID:BDSC_34775 |
| cno | canoe | BDSC | 33367 | RRID:BDSC_33367 |
| Cka | Connector of kinase to AP-1 | BDSC | 34522 | RRID:BDSC_34522 |
| csw | corkscrew | BDSC | 33619 | RRID:BDSC_33619 |
| Cdk8 | Cyclin-dependent kinase 8 | BDSC | 35324 | RRID:BDSC_35324 |
| Delta | Delta | BDSC | 34322 | RRID:BDSC_34322 |
| dlg1 | discs large 1 | BDSC | 36771 | RRID:BDSC_36771 |
| dl | dorsal | BDSC | 32934 | RRID:BDSC_32934 |
| γTub37C | Gamma Tubulin at 37C | BDSC | 32513 | RRID:BDSC_32513 |
| glo | glorund | BDSC | 33668 | RRID:BDSC_33668 |
| hop | hopscoth | BDSC | 32966 | RRID:BDSC_32966 |
| key | kenny | BDSC | 35572 | RRID:BDSC_35572 |
| mTor | Megator | BDSC | 32941 | RRID:BDSC_32941 |
| mthl1 | methuselah-like 1 | BDSC | 57699 | RRID:BDSC_57699 |
| mri | mrityu | BDSC | 35165 | RRID:BDSC_35165 |
| nanos | nanos | BDSC | 33973 | RRID:BDSC_33973 |
| okr | okra | BDSC | 33707 | RRID:BDSC_33707 |
| Pka-C1 | Protein kinase, cAMP-dependent, catalytic subunit 1 | BDSC | 35169 | RRID:BDSC_35169 |
| put | punt | BDSC | 39025 | RRID:BDSC_39025 |
| Raf | Raf oncogene | BDSC | 41589 | RRID:BDSC_41589 |
| rhi | rhino | BDSC | 34071 | RRID:BDSC_34071 |
| rl | rolled | BDSC | 34855 | RRID:BDSC_34855 |
| SAK | SAK kinase | BDSC | 57221 | RRID:BDSC_57221 |
| sax | saxophone | BDSC | 55865 | RRID:BDSC_55865 |
| skd | skuld | BDSC | 34630 | RRID:BDSC_34630 |
| sle | slender lobes | BDSC | 32969 | RRID:BDSC_32969 |
| slpr | slipper | BDSC | 32948 | RRID:BDSC_32948 |
| Smox | Smad on X | BDSC | 41670 | RRID:BDSC_41670 |
| SmD1 | Small ribonucleoprotein particle protein | BDSC | 34834 | RRID:BDSC_34834 |
| Sos | Son of sevenless | BDSC | 34833 | RRID:BDSC_34833 |
| spz | spatzle | BDSC | 28538 | RRID:BDSC_28538 |
| spn-E | spindle-E | BDSC | 34808 | RRID:BDSC_34808 |
| spen | split ends | BDSC | 33398 | RRID:BDSC_33398 |
| tkv | thickveins | BDSC | 40937 | RRID:BDSC_40937 |
| Trxr1 | Thioredoxin reductase 1 | BDSC | 32984 | RRID:BDSC_32984 |
| tj-Gal4 | traffic jam Gal4 | Gontijo Lab | NA | NA |
| Tl | Toll | BDSC | 35628 | RRID:BDSC_35628 |
| tor | Torso | BDSC | 33627 | RRID:BDSC_33627 |
| tsl | torso-like | BDSC | 56967 | RRID:BDSC_56967 |
| tub | tube | BDSC | 66960 | RRID:BDSC_66960 |
| Vps15 | Vacuolar protein sorting 15 | BDSC | 35209 | RRID:BDSC_35209 |
| wit | wishful thinking | BDSC | 41906 | RRID:BDSC_41906 |
| dilp8 | Drosophila Insulin-like peptide 8 | VDRC | 102604 | NA |

**Supplementary Table 2 – Composition of yeast based medium (YBM).**

| | Ingredient | Quantity | Ingredient References |
|---|---|---|---|
| **Yeast Based Medium (YBM) recipe (1L)** | Barley Malt Syrup | 80 g | Provida, G109115B |
| | Beetroot Syrup | 22 g | Grafschafter Krautfabrik Josef Schmitz |
| | Biological Corn Flour | 80 g | Farinhas Paulino Horta |
| | Instant Yeast | 18 g | Mauripan, ICOPA |
| | Soy Flour | 10 g | A CENTAZZI, LDA |
| | Agar | 8 g | MB14801, NZYTech |
| | Nipagin 15% in EtOH | 12 mL | 25605.293, Avantor-VWR. |
| | Propionic Acid >99% | 8 mL | ACROS ORGANICS, 149300025. |
| | Water (% varies) | 1.1-1.25 L | |

### Supplementary Table 3 – Composition of holidic media (HM).

| | HM | HM - sucrose | HM - non essential amino acids | HM - essential amino acids | HM - cholesterol | HM - vitamins | HM - metal ions | HM - nucleic acids | HM - lipids | HM - folic acid |
|---|---|---|---|---|---|---|---|---|---|---|
| **Referred to as** | **Complete HM** | **w/o sucrose** | **w/o NEAA** | **w/o EAA** | **w/o cholesterol** | **w/o vitamins** | **w/o metal ions** | **w/o nucleic acids** | **w/o lipids** | **w/o folic acid** |
| Essential amino acids | 60.51 mL | 60.51 mL | 116.78 mL (a) | 00.00 mL | 60.51 mL | 60.51 mL | 60.51 mL | 60.51 mL | 60.51 mL | 60.51 mL |
| L-isoleucine | 1.12 g | 1.12 g | 2.16 g (a) | 00.00 mL | 1.12 g | 1.12 g | 1.12 g | 1.12 g | 1.12 g | 1.12 g |
| L-leucine | 2.03 g | 2.03 g | 3.92 g (a) | 00.00 mL | 2.03 g | 2.03 g | 2.03 g | 2.03 g | 2.03 g | 2.03 g |
| Non-essential amino acids | 60.51 mL | 60.51 mL | 0.00 mL | 113.76 mL (a) | 60.51 mL | 60.51 mL | 60.51 mL | 60.51 mL | 60.51 mL | 60.51 mL |
| L-glutamate | 15.19 mL | 15.19 mL | 29.32 mL (a) | 28.56 mL (a) | 15.19 mL | 15.19 mL | 15.19 mL | 15.19 mL | 15.19 mL | 15.19 mL |
| L-tyrosine | 0.93 g | 0.93 g | 0.00 g | 1.75 g (a) | 0.93 g | 0.93 g | 0.93 g | 0.93 g | 0.93 g | 0.93 g |
| L-cysteine (HCl) | 6.83 mL | 6.83 mL | 0.00 g | 12.84 g | 6.83 mL | 6.83 mL | 6.83 mL | 6.83 mL | 6.83 mL | 6.83 mL |
| Cholesterol | 15.00 mL | 15.00 mL | 15.00 mL | 15.00 mL | 00.00 mL | 15.00 mL | 15.00 mL | 15.00 mL | 15.00 mL | 15.00 mL |
| CaCl2 | 1.00 mL | 1.00 mL | 1.00 mL | 1.00 mL | 1.00 mL | 1.00 mL | 00.00 mL | 1.00 mL | 1.00 mL | 1.00 mL |
| MgSO4 | 1.00mL | 1.00mL | 1.00mL | 1.00mL | 1.00mL | 1.00mL | 00.00 mL | 1.00mL | 1.00mL | 1.00mL |
| CuSO4 | 1.00 mL | 1.00 mL | 1.00 mL | 1.00 mL | 1.00 mL | 1.00 mL | 00.00 mL | 1.00 mL | 1.00 mL | 1.00 mL |
| FeSO4 | 1.00mL | 1.00mL | 1.00mL | 1.00mL | 1.00mL | 1.00mL | 00.00 mL | 1.00mL | 1.00mL | 1.00mL |
| MnCl2 | 1.00 mL | 1.00 mL | 1.00 mL | 1.00 mL | 1.00 mL | 1.00 mL | 00.00 mL | 1.00 mL | 1.00 mL | 1.00 mL |
| ZnSO4 | 1.00mL | 1.00mL | 1.00mL | 1.00mL | 1.00mL | 1.00mL | 00.00 mL | 1.00mL | 1.00mL | 1.00mL |
| Nucleic acids & Lipids | 8.00 mL | 8.00 mL | 8.00 mL | 8.00 mL | 8.00 mL | 8.00 mL | 8.00 mL | 8.00 mL (Lipid solution) | 8.00 mL (NA solution) | 8.00 mL |
| Vitamins | 21.00 mL | 21.00 mL | 21.00 mL | 21.00 mL | 21.00 mL | 00.00 mL | 21.00 mL | 21.00 mL | 21.00 mL | 21.00 mL |
| Folic acid | 1.00 mL | 1.00 mL | 1.00 mL | 1.00 mL | 1.00 mL | 1.00 mL | 1.00 mL | 1.00 mL | 1.00 mL | 00.00 mL |
| Acetic acid buffer | 100.00 mL | 100.00 mL | 100.00 mL | 100.00 mL | 100.00 mL | 100.00 mL | 100.00 mL | 100.00 mL | 100.00 mL | 100.00 mL |
| Sucrose | 17.12 g | 0.00 g | 17.12 g | 17.12 g | 17.12 g | 17.12 g | 17.12 g | 17.12 g | 17.12 g | 17.12 g |
| Agar | 20.00 g | 20.00 g | 20.00 g | 20.00 g | 20.00 g | 20.00 g | 20.00 g | 20.00 g | 20.00 g | 20.00 g |
| Nipagin | 15.00 mL | 15.00 mL | 15.00 mL | 15.00 mL | 15.00 mL | 15.00 mL | 15.00 mL | 15.00 mL | 15.00 mL | 15.00 mL |
| Propionic Acid | 6.00 mL | 6.00 mL | 6.00 mL | 6.00 mL | 6.00 mL | 6.00 mL | 6.00 mL | 6.00 mL | 6.00 mL | 6.00 mL |
| Milli Q H2O | Adjust to 1 L | Adjust to 1 L | Adjust to 1 L | Adjust to 1 L | Adjust to 1 L | Adjust to 1 L | Adjust to 1 L | Adjust to 1 L | Adjust to 1 L | Adjust to 1 L |

(a) The amount of these nutrients was increased to adjust concentration of biological active nitrogen to 200 mM as in the complete HM. In diets where neAAs were removed, L-glutamate was still added.

**Supplementary Table 4 – Composition of apple juice plates.**

| | Ingredient | Quantity | Ingredient References |
|---|---|---|---|
| **Apple juice agar plates recipe** | Agar | 19.5 g | MB14801, NZYTech |
| | Sugar | 20 g | Açúcar Branco de Cana Sidul |
| | Apple Juice | 250 mL | Compal 100% Maçã, 100% Fruta. |
| | Nipagin 10% in EtOH | 10 mL | 25605.293, Avantor-VWR. |
| | Water | 750 mL | |

**Supplementary Table 5 – Primers used for RT-qPCR.**

| Primer Name | Associated gene | Associated gene (CG number) | Sequence (5´-3') |
|---|---|---|---|
| dilp8 Fw | dilp8 | CG14059 | CGACAGAAGGTCCATCGAGT |
| dilp8 Rv | dilp8 | CG14059 | GATGCTTGTTGTGCGTTTTG |
| Act42A Fw | Act42A | CG12051 | CAGGCGGTGCTTTCTCTCTA |
| Act42A Rv | Act42A | CG12051 | AGCTGTAACCGCGCTCAGTA |
| Rp49 Fw | Rp49 | CG7939 | GCACTCTCTGTTGTCGATACCCTTG |
| Rp49 Rv | Rp49 | CG7939 | AGCGCACCAAGCACTTCATC |

### Supplementary Table 6 – p-values of fertility measurements.

| | # eggs/female | | | fraction of viable eggs | | | estimated progeny | | |
|---|---|---|---|---|---|---|---|---|---|
| **Gene name** | **Phenotype** | **p-value** | **p-value** | **Phenotype** | **p-value** | **p-value** | **Phenotype** | **p-value** | **p-value** |
| *mri* | decrease | *** | 0.0009 | decrease | * | 0.0142 | decrease | ** | 0.004 |
| *Vps15* | decrease | *** | 0.0002 | – | ns | 0.7768 | decrease | * | 0.0162 |
| *Argk1* | – | ns | 0.3298 | decrease | ** | 0.0024 | decrease | ** | 0.002 |
| *Tl* | decrease | ** | 0.0076 | – | ns | >0.9999 | decrease | * | 0.0327 |
| *mthl1* | – | ns | >0.9999 | – | ns | >0.9999 | – | ns | >0.9999 |
| *spz* | – | ns | 0.5658 | decrease | **** | <0.0001 | decrease | ** | 0.0058 |
| *γTub37C* | – | ns | >0.9999 | – | ns | >0.9999 | – | ns | >0.9999 |
| *spen* | – | ns | 0.347 | – | ns | 0.0834 | – | ns | >0.9999 |
| *csw* | decrease | **** | <0.0001 | – | ns | 0.0506 | decrease | * | 0.037 |
| *slpr* | – | ns | 0.2273 | – | ns | >0.9999 | – | ns | 0.542 |
| *tor* | – | ns | 0.0981 | decrease | ** | 0.0033 | – | ns | >0.9999 |
| *okr* | – | ns | >0.9999 | – | ns | 0.1921 | – | ns | >0.9999 |
| *bcd* | decrease | * | 0.0432 | decrease | ** | 0.0073 | decrease | * | 0.0136 |
| *nanos* | decrease | ** | 0.0023 | decrease | **** | <0.0001 | decrease | *** | 0.0003 |
| *dl* | decrease | *** | 0.0007 | decrease | *** | 0.0005 | decrease | *** | 0.0002 |
| *mTor* | decrease | * | 0.0305 | decrease | **** | <0.0001 | decrease | ** | 0.0019 |
| *sle* | decrease | **** | <0.0001 | decrease | *** | 0.0007 | decrease | **** | <0.0001 |
| *cno* | – | ns | >0.9999 | – | ns | 0.1184 | – | ns | >0.9999 |
| *glo* | – | ns | >0.9999 | – | ns | 0.7338 | – | ns | 0.6887 |
| *Delta* | decrease | * | 0.0403 | – | ns | 0.0642 | decrease | ** | 0.0068 |
| *Cka* | – | ns | 0.186 | – | ns | 0.3912 | – | ns | >0.9999 |
| *cact* | – | ns | 0.2163 | decrease | **** | <0.0001 | decrease | ** | 0.0068 |
| *smD1* | – | ns | >0.9999 | – | ns | 0.3896 | – | ns | >0.9999 |
| *rl* | – | ns | 0.7992 | decrease | **** | <0.0001 | – | ns | 0.1819 |
| *sax* | increase | *** | 0.0005 | – | ns | >0.9999 | – | ns | 0.4042 |
| *Trxr1* | – | ns | 0.0958 | – | ns | >0.9999 | – | ns | >0.9999 |
| *skd* | decrease | ** | 0.0067 | decrease | ** | 0.0034 | – | ns | 0.0798 |
| *spn-E* | decrease | **** | <0.0001 | – | ns | >0.9999 | decrease | * | 0.0157 |
| *dlg1* | decrease | *** | 0.0004 | – | ns | 0.178 | – | ns | >0.9999 |
| *rhi* | increase | *** | 0.0005 | – | ns | >0.9999 | – | ns | 0.6353 |
| *aPKC* | – | ns | 0.0961 | decrease | ** | 0.0033 | – | ns | 0.0641 |
| *put* | – | ns | 0.4183 | – | ns | >0.9999 | – | ns | 0.5066 |
| *Smox* | decrease | **** | <0.0001 | – | ns | >0.9999 | decrease | *** | 0.0005 |
| *tsl* | – | ns | 0.5608 | increase | * | 0.0449 | – | ns | 0.9158 |
| *tub* | – | ns | 0.9995 | – | ns | >0.9999 | – | ns | 0.9925 |
| *tkv* | **NA** | **NA** | **NA** | **NA** | **NA** | **NA** | **NA** | **NA** | **NA** |
| *SAK* | **NA** | **NA** | **NA** | **NA** | **NA** | **NA** | **NA** | **NA** | **NA** |
| *babo* | – | ns | 0.0597 | – | ns | >0.9999 | decrease | * | 0.0182 |
| *Pka-C1* | decrease | **** | <0.0001 | decrease | *** | 0.0005 | decrease | **** | <0.0001 |
| *Cdk8* | decrease | **** | <0.0001 | decrease | ** | 0.0071 | decrease | **** | <0.0001 |
| *key* | decrease | * | 0.0313 | – | ns | 0.8137 | decrease | ** | 0.0027 |
| *Raf* | – | ns | 0.5723 | – | ns | 0.1419 | – | ns | 0.7978 |
| *hop* | – | ns | 0.1505 | decrease | **** | <0.0001 | decrease | **** | <0.0001 |
| *Sos* | **NA** | **NA** | **NA** | **NA** | **NA** | **NA** | **NA** | **NA** | **NA** |
| *wit* | – | ns | 0.1185 | – | ns | 0.9984 | decrease | *** | 0.0003 |

### Supplementary Table 7 – p-values of behavior measurements.

| | # sips on yeast | | | # sips on sucrose | | |
|---|---|---|---|---|---|---|
| **Gene name** | **Phenotype** | **p-value** | **p-value** | **Phenotype** | **p-value** | **p-value** |
| *mri* | increase | * | p=0.0222 | decrease | *** | 0.0003 |
| *Vps15* | – | ns | p=0.0841 | decrease | ** | 0.0062 |
| *Argk1* | increase | *** | p=0.0002 | – | ns | 0.1909 |
| *Tl* | – | ns | p=0.0525 | decrease | * | 0.0103 |
| *mthl1* | increase | * | p=0.0373 | – | ns | >0.9999 |
| *spz* | increase | *** | p=0.0006 | – | ns | >0.9999 |
| *γTub37C* | increase | ** | p=0.0038 | – | ns | 0.08 |
| *spen* | – | ns | p=0.6112 | increase | **** | <0.0001 |
| *csw* | – | ns | p=0.2684 | – | ns | 0.0584 |
| *slpr* | – | ns | p=0.9689 | increase | **** | <0.0001 |
| *tor* | – | ns | p=0.5602 | increase | **** | <0.0001 |
| *okr* | increase | * | p=0.0499 | – | ns | 0.069 |
| *bcd* | increase | **** | <0.0001 | – | ns | >0.9999 |
| *nanos* | – | ns | p=0.1516 | – | ns | >0.9999 |
| *dl* | increase | * | p=0.0147 | – | ns | >0.9999 |
| *mTor* | – | ns | >0.9999 | – | ns | 0.5456 |
| *sle* | increase | ** | p=0.0023 | – | ns | >0.9999 |
| *cno* | – | ns | p=0.2809 | increase | * | 0.0313 |
| *glo* | – | ns | p=0.9359 | increase | **** | <0.0001 |
| *Delta* | – | ns | >0.9999 | – | ns | 0.67 |
| *Cka* | – | ns | >0.9999 | – | ns | 0.5273 |
| *cact* | increase | * | p=0.0114 | – | ns | >0.9999 |
| *smD1* | decrease | ** | p=0.0027 | – | ns | >0.9999 |
| *rl* | increase | * | p=0.0128 | – | ns | >0.9999 |
| *sax* | decrease | * | p=0.0275 | – | ns | >0.9999 |
| *Trxr1* | – | ns | >0.9999 | – | ns | >0.9999 |
| *skd* | – | ns | >0.9999 | – | ns | >0.9999 |
| *spn-E* | decrease | *** | p=0.0003 | decrease | * | 0.0232 |
| *dlg1* | – | ns | >0.9999 | – | ns | 0.4954 |
| *rhi* | – | ns | p=0.2524 | increase | * | 0.0417 |
| *aPKC* | increase | ** | p=0.0024 | increase | * | 0.0212 |
| *put* | increase | **** | <0.0001 | – | ns | 0.3434 |
| *Smox* | increase | * | p=0.0377 | increase | *** | 0.0003 |
| *tsl* | – | ns | >0.9999 | increase | ** | 0.0036 |
| *tub* | – | ns | p=0.1159 | increase | ** | 0.0053 |
| *tkv* | – | ns | >0.9999 | – | ns | >0.9999 |
| *SAK* | – | ns | >0.9999 | – | ns | 0.1063 |
| *babo* | increase | ** | p=0.0013 | – | ns | >0.9999 |
| *Pka-C1* | increase | ** | p=0.0013 | – | ns | >0.9999 |
| *Cdk8* | – | ns | p=0.6348 | – | ns | 0.0558 |
| *key* | – | ns | >0.9999 | – | ns | >0.9999 |
| *Raf* | – | * | 0.0438 | – | ns | 0.8241 |
| *hop* | increase | **** | <0.0001 | – | ns | >0.9999 |
| *Sos* | – | ns | p=0.0508 | decrease | *** | 0.0005 |
| *wit* | increase | ** | p=0.0013 | – | ns | >0.9999 |
